# A Hybrid Knowledge- and Data-driven Model for Automatic Assessment of Induced Spiking Patterns in C-fiber Microneurography: Methodological Development

**DOI:** 10.1101/2025.10.13.681964

**Authors:** Alina Troglio, Andrea Fiebig, Anna Maxion, Ekaterina Kutafina, Barbara Namer

**Affiliations:** Institute of Neurophysiology, RWTH Aachen University Hospital, Aachen, Germany; Department of Anesthesiology, Intensive Care, Emergency and Pain Medicine, University Hospital Würzburg, Center for Interdisciplinary Pain Medicine, Würzburg, Germany; Research Group Neuroscience, Interdisciplinary Centre for Clinical Research (IZKF), Faculty of Medicine, RWTH Aachen University, Aachen, Germany; Institute for Computational Biomedicine, RWTH Aachen University, Aachen, Germany; Institute for Biomedical Informatics, Faculty of Medicine, University Hospital Cologne, University of Cologne, Cologne, Germany

**Keywords:** microneurography, spike detection, spike sorting, machine learning, pain, itch, C-fiber

## Abstract

Analyzing temporal spike patterns in C-fibers recorded via microneurography is challenging due to the use of a single recording electrode, waveform variability, and high similarity of spike shapes across neurons, limiting the interpretation of sensory coding, such as pain and itch. We present a computational pipeline combining peak detection and supervised classification for spike sorting to improve the analysis of discharges, identified through activity-dependent conduction velocity changes. In the knowledge-driven step, we extract spike templates from electrically evoked spikes obtained during low-frequency stimulation and focus on the “best” template as the fiber of interest. Spike detection is further restricted to intervals showing activity-dependent latency shifts, substantially reducing the search space compared to unsupervised clustering. In the data-driven steps, we systematically evaluate three feature sets and machine learning models: One-class SVM, SVM, and XGBoost. For the evaluation, we created a specialized stimulation protocol, providing reliable ground truth labels for all electrically evoked spikes, allowing precise spike time-locking. Compared to Spike2 software, our approach achieved higher F1-scores and reduced false positives, indicating improved spike sorting. Although XGBoost achieved the highest median F1-scores, optimal performance was dependent on individual combinations of feature sets and models for each recording. In some recordings with many nerve fibers and a low signal-to-noise ratio, reliable sorting was not feasible. This highlights the necessity to determine sortability and optimal configurations for individual recordings. To illustrate the potential of our approach to sensory spike train analysis, we present a proof-of-concept application of the pipeline to chemically induced C-fiber activity. These findings represent an important step toward reliable analysis of activity associated with pain and itch signaling.

## Introduction

Sensory perception in the peripheral nervous system depends not only on the presence or absence of action potentials (spikes) but also on the precise temporal patterns in which they are fired [1, 2]. Spike patterns defined by firing regularity, bursting, and adaptation carry information about stimulus quality and intensity [3]. In the central nervous system, the temporal dynamics of neural activity are known to shape synaptic strength and influence network plasticity [4]. Moreover, spikes in peripheral afferents can modulate the behavior of extraneural cells, such as immune or glial cells, through activity-dependent neuroimmune interactions [5]. These insights suggest that spike timing and patterns in peripheral neurons are not merely epiphenomena but may play a critical role in encoding and modulating sensations, particularly in complex modalities like pain and itch.

Despite this, the spike patterns of peripheral nociceptors and pruriceptors, especially in humans, have received relatively little attention. Most single-electrode recordings of peripheral nerve fibers in human (microneurography) studies have focused on indirect assessment of spike rates or thresholds rather than temporal structure, leaving a major gap in our understanding of how these neurons convey nuanced sensory information [6]. This analysis is especially relevant in pathological conditions such as neuropathic pain, as some primary afferent fibers exhibit spontaneous, continuous, and irregular firing patterns that are not evoked by external stimuli [7]. These complex firing patterns may have functional consequences for downstream processing and symptom manifestation. For more general neuroscientific questions of sensory encodings of itch and pain, it must be noted that the same nerve fibers are able to encode itchy and painful heat stimuli [8, 9]. In our previous work, current temporal discharge pattern theories propose that pruritogen-induced stimulation evokes specific burst-like firing with gaps between activity bursts, whereas stimuli related to the sensation of pain typically induce more continuous discharges followed by gradual adaptation [8]. The bursting is characterized by discharges for approximately 4-12 seconds and silent periods of approximately 20-120 seconds. However, without reliable spike pattern analysis, those hypotheses and questions cannot be explored and answered.

Microneurography enables direct extracellular recordings from single afferent fibers in awake humans, offering insights into the dynamics of sensory encoding at the level of individual neurons [10]. However, because single tungsten microelectrodes capture activity from multiple nearby fibers, spike detection and sorting are essential before any meaningful analysis of spike patterns can be performed. This process is particularly difficult in peripheral nerve recordings, especially for unmyelinated nerve fibers, which are also abundantly present in the spinal cord and brain. In these recordings, spike amplitudes are small with typically low signal-to-noise ratios, conduction velocities are activity-dependent, and waveform shapes are very similar between different fibers and variable in a single nerve fiber [11, 12]. In addition to these challenges, microneurography experiments may exhibit slow waveform drift caused by long recording durations. Since such drift can influence spike sorting, monitoring waveform stability throughout the experiment is essential in microneurography.

Numerous spike sorting algorithms have been developed, ranging from traditional clustering techniques to algorithms such as Kilosort [13], MountainSort [14], and WaveClus [15], as well as benchmarking and comparison frameworks such as SpikeInterface [16] and SpikeForest [17]. However, most of these algorithms and tools are optimized for high-density or multi-site electrode arrays [18]. They often rely on spatial separation of units, timing effects, or stable waveform shapes across channels, assumptions that do not hold in single-electrode microneurography. In our previous work, we showed that unsupervised spike sorting methods have limited reliability when applied to single-channel microneurography recordings [19]. Since microneurography experiments typically include periodic low-frequency electrical stimulation, a subset of spikes can be indirectly labeled using the marking method [20]. In the marking method, this stimulation ensures that each recording contains a reliable set of electrically evoked spikes with known fiber labels. In our earlier feasibility analysis, these consistently available labeled spikes were used as training data for a supervised spike-sorting approach [19].

In this manuscript, we present an open-source spike detection and sorting pipeline for noisy single-electrode recordings in peripheral nerves, focusing on C-fiber nociceptors relevant to pain and itch. Our approach is optimized for recordings with low-amplitude spikes and builds on our earlier findings showing that spikes evoked by regular low-frequency stimuli provide training data for supervised classifiers with different feature sets [19]. Here, we introduce two key extensions. First, we designed a specialized stimulation protocol that enhances the traditional marking method [20], establishing reliable ground truth data for all electrically evoked spikes. Second, we developed an improved computational pipeline combining peak detection with supervised classification, utilizing latency and activity-dependent conduction velocity changes to constrain and refine spike detection. We benchmark our methods against Spike2, which allows spike sorting by human expert template creation and is routinely used in the microneurography community [21, 22], using ground truth data. To illustrate real-world applicability, we applied our pipeline to a recording with pruritogen (itch-inducing) injection, evoking spikes in C-fibers. By combining experimental ground truth data with supervised classification, this work provides a foundation for decoding peripheral nerve activity and advancing the study of sensory coding.

## Methods

### Data and ethical approval

A continuous microneurography study was conducted between 01/11/2019-31/12/2024 at the RWTH Aachen University Medical Faculty (Namer’s Lab), approved by the Ethics Board of the University Hospital RWTH Aachen with numbers Vo-Nr. EK141-19 and Vo-Nr. EK143- 21. The participants were comprehensively informed about the experimental procedure, provided their written informed consent, and the study was conducted according to the Declaration of Helsinki. Microneurography recordings were acquired during laboratory-based experiments designed to investigate peripheral nerve activity and to develop and evaluate signal processing and analysis methods. In this research, we analyze the data from seven healthy volunteers (one male, six females). None of the participants had any neurological, dermatological, or other chronic medical conditions, and none had taken regular medication or any acute medication within 24 hours prior to the experiments. Six recordings include fully available ground truth labels for all electrically evoked spikes, and one recording with chemical stimulation is included to demonstrate a proof-of-concept application.

### Experimental setup

With microneurography, spikes from individual C-fibers within the cutaneous C-fiber fascicles of the superficial peroneal nerve were recorded. A detailed explanation of this technique can be found in [8, 11]. For the experiments, a recording electrode (Frederick-Haer, Bowdoinham, ME, USA) was positioned near the nerve bundle, and innervation territories were mapped using a 0.5-diameter tipped electrode. Once the targeted territories were identified, 0.2 mm diameter needle electrodes (Frederick-Haer) were inserted into the skin for intracutaneous electrical stimulation via a Digitimer DS7 constant current stimulator. Neural signals were amplified and processed in parallel using a Neuro Amp EX (ADInstruments) amplifier, followed by an additional bandpass filter (500-1000 Hz) and a 50 Hz notch filter to minimize electrical noise. The data acquisition was performed with two systems in parallel: 1) with a sampling rate of 10 kHz using an analog-to-digital converter from National Instruments and the software Dapsys (www.dapsys.net) by Brian Turnquist [12], and 2) Power1401 as a digital analogue converter from Cambridge Electronics Design and the software Spike2 v10.08.

### Marking method

In microneurography, the marking method is routinely used as a standard approach to differentiate individual C-fiber responses and C-fiber types [11, 20]. After identifying the receptive field and determining the electrical threshold required to reliably activate the recorded fibers, low-frequency “background” electrical stimuli are applied. These background stimuli evoke exactly one spike per activated fiber and are delivered at a fixed interval (here, every 4 seconds), creating a predictable time window in which the evoked spike is expected due to a constant conduction velocity of C-fibers under low-frequency stimulation. The signal is segmented relative to each background stimulus onset, allowing spikes originating from a single C-fiber to align temporally at a consistent latency. The microneurography recording is visualized in a raster-like “waterfall plot”, which creates temporal alignment of spikes regardless of amplitude (see Figure 1a, blue and green circles). These vertically aligned, repeated spikes are referred to as *tracks*.

**Figure 1.**
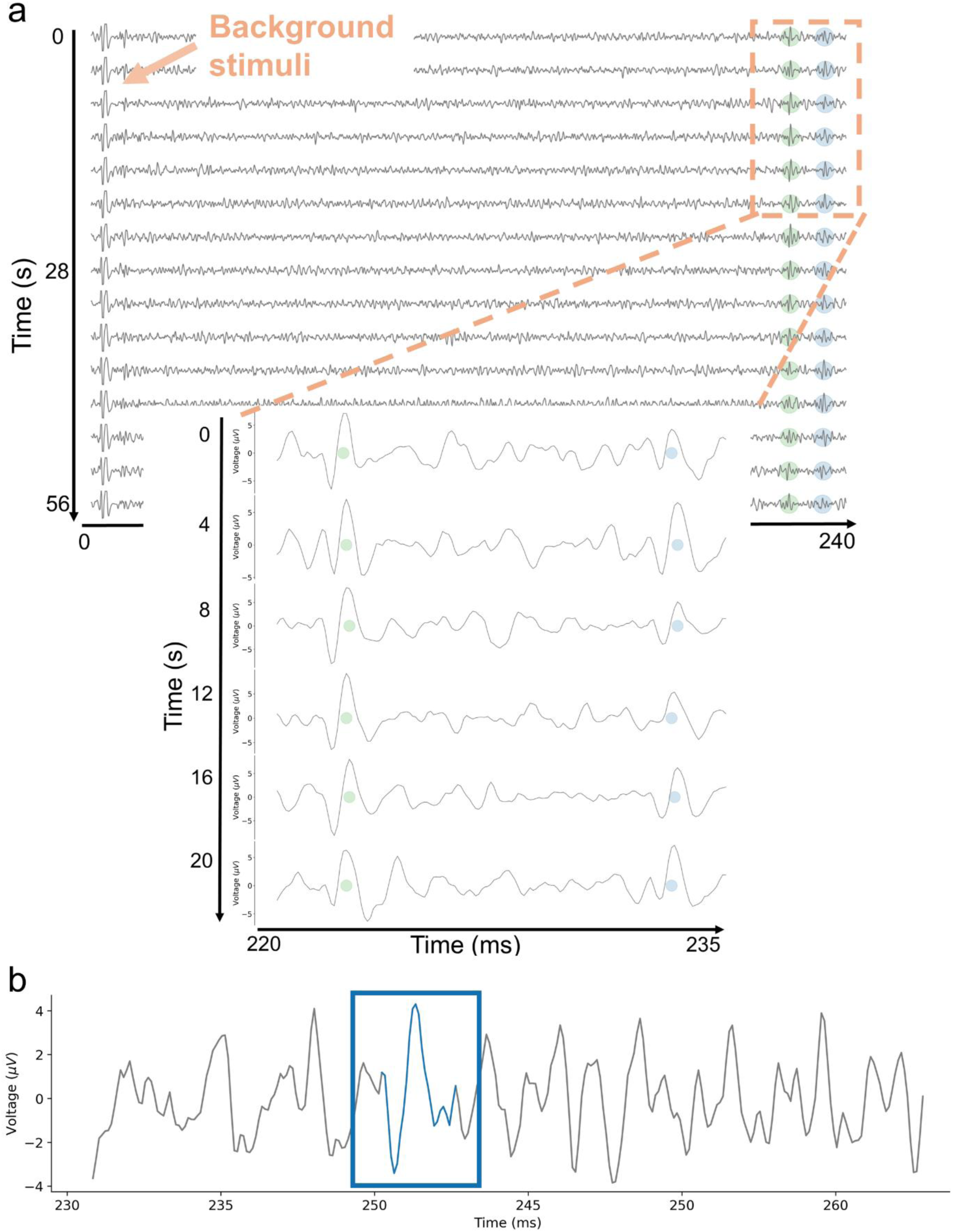
**(a)** Example waterfall plot of two tracks. The signal is segmented every 4 seconds, beginning with the electrical background stimulus. Two tracks are visible, marked by the green and blue dots, indicating the activity of two C-fibers. The conduction velocity stays almost constant, leading to vertical alignment of the elicited spikes. **(b)** An example spike at the noise level. The spike in the blue box would not be detectable without the vertical alignment of repeated responses enabled by the marking method and waterfall representation. This temporal alignment facilitates the identification of low-amplitude spikes by leveraging consistent latencies evoked by background stimuli.

The resulting tracks enable reliable identification and sorting with tracking algorithms [12], even for spikes near the noise level (see Figure 1b, blue rectangle). Since the track reflects consistent latency across many repetitions, it remains robust to individual missed spikes, rather than being dependent on a single response. When a fiber is activated shortly before an electrical background pulse, its conduction velocity temporarily decreases, leading to a delayed response to the subsequent background pulse (see Figure 2a, line 4). This latency shift, which scales approximately with the number of preceding spikes, is known as activity-dependent conduction velocity slowing (ADS) [23]. ADS serves as a functional “marker” for individual fibers and allows for semi-quantitative estimation of fiber activity.

**Figure 2.**
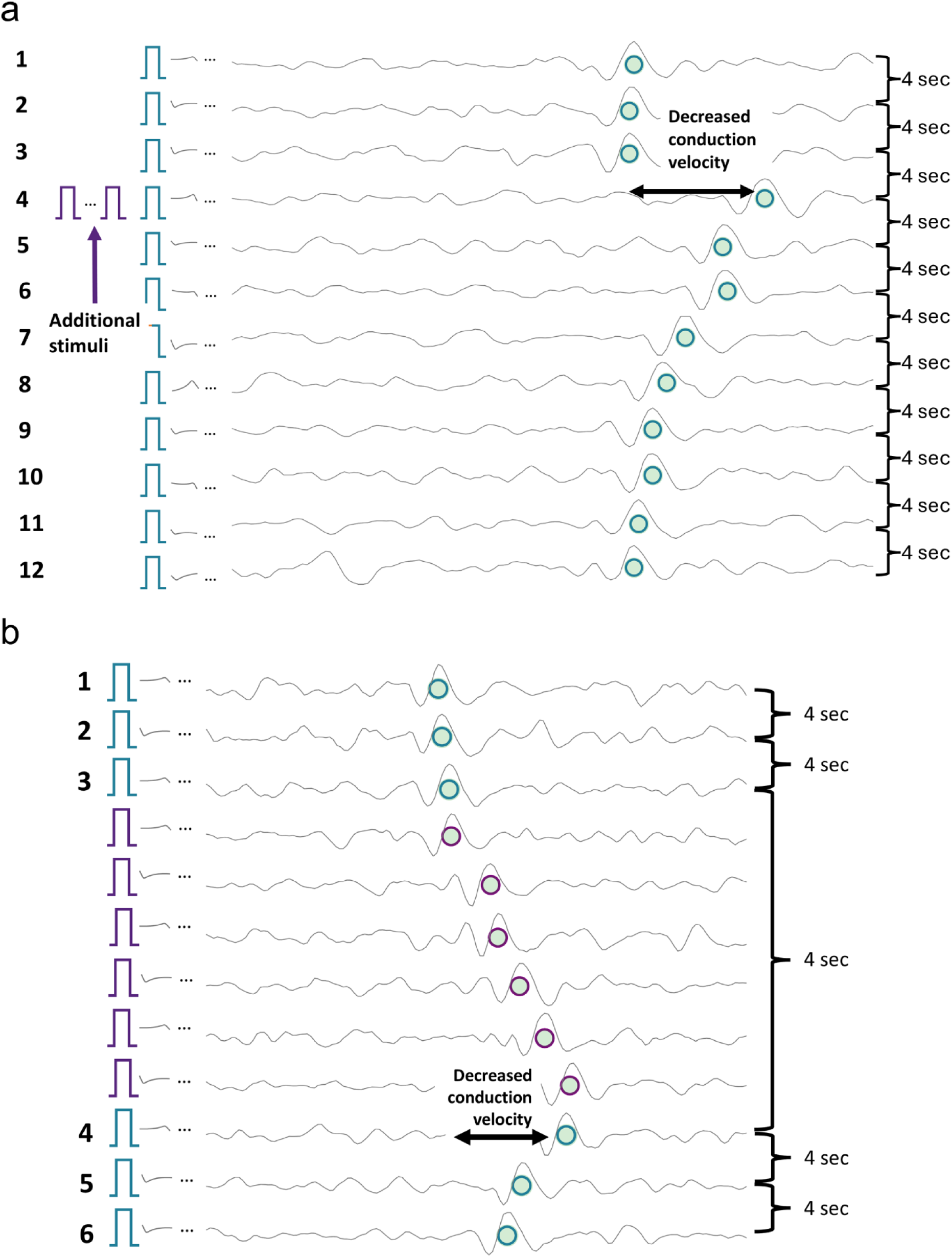
(a) Example of activity-dependent slowing with the marking method and additional stimuli. Each line begins with the periodic background pulse indicated by the blue rectangle, resulting in signal segments with a duration of 4 seconds. When no extra activity is observed (lines 1-3), the spikes (green dots) align vertically due to the almost constant conduction velocity. When additional (electrical) stimuli are applied, there is a decrease in conduction velocity (line 4). Responses to background stimuli are marked by green dots with a blue circle. (b) Example of activity-dependent slowing stimulated with the ground truth protocol. As in the routinely used marking method, there are background stimuli every 4 seconds and additional pulses as in (a). In contrast to the “normal” marking method, the signal is cut before each stimulation pulse, whether it is a background pulse or an additional pulse. This ensures that all spikes are aligned on the green track and not only the ones evoked by background stimuli. Responses to background stimuli remain marked by green dots with blue circles, while responses to additional stimuli are indicated by purple rectangles and circles, both for the stimuli and the evoked spikes.

The marking method reliably differentiates spikes from individual C-fibers by assigning them to vertical tracks formed by responses time-locked to low-frequency stimulation. These tracks are routinely present and can support training of supervised machine learning, but their limited number motivates the use of lightweight algorithms. In typical experimental protocols, this background stimulation is combined with additional stimuli applied in parallel to mimic fiber activity and evoked sensations like itch and pain [8, 11]. However, this spike identification and sorting are inherently limited to spikes that occur due to background pulses. Spikes evoked independently of this background stimulation, such as electrically, mechanically, spontaneously or pruritogen-evoked activity, do not align with these tracks and therefore cannot be directly detected or assigned to specific fibers. Their influence is only indirectly observable through activity-dependent conduction-velocity slowing.

To overcome this limitation without altering the foundation of the marking method, we implemented a hardware- and software-level extension of the stimulation protocol that preserves the background stimuli but redefines how additional electrical stimuli are processed. The goal is to create recordings in which all electrically evoked spikes, including those mimicking extra fiber activity, are time-locked and reliably traceable.

### “Ground truth” stimulation protocol for model development and validation

In our developed “ground truth” protocol, the stimulation pattern remains identical to other electrical experiments: periodic 0.25 Hz background stimuli with additional pulses inserted between background pulses to mimic extra fiber activity. The key modification is the hardware/software interpretation of these events. In contrast to the routinely used marking method, the ground truth protocol is an engineered extension used solely for algorithm training and validation. It differs fundamentally in that all stimuli, including the inserted additional pulses, are treated as background stimuli, meaning that each stimulus defines its own segmentation window and allows any evoked spike to be aligned to the corresponding fiber track.

As a result, the segmentation windows in the waterfall representation are dynamically cut after each stimulus, producing variable-length intervals between consecutive stimuli rather than the fixed 4s windows used in the standard marking method (see Figure 2b). This ensures that spikes occurring between two classical background pulses are projected onto the vertical tracks and thus become reliably detectable and sortable.

Importantly, as in the marking method, the approach is robust to occasionally missed spikes. A fiber may fail to respond to an additional pulse, but this would only result in a smaller or absent latency shift and does not compromise the interpretability of the track. The essential aim of the protocol is to ensure that whenever a spike is electrically evoked, it is time-locked and can be correctly labeled.

Together, these modifications generate recordings with fully verified spike timestamps and fiber identities, providing a reliable ground truth for validating spike detection and sorting algorithms under realistic single-electrode noise conditions.

### Additional pulse parameters in the ground truth protocol

The number and timing of additional pulses in the ground truth protocol were not fixed but varied across recordings. The extended marking method approach is independent of the exact number of inserted pulses, because each stimulus is processed identically by the hardware/software and provides a time-locked response for fiber-specific spike assignment. Whether an interval contains two or ten pulses, the resulting latency tracks remain continuous, stable, and fully interpretable. In Table 1, the range of additional pulses, the inter-stimulus frequency, and the distribution of distances to the next background stimulus are summarized. These values highlight the diversity of stimulation patterns and show that it is the technical changes of the protocol, not any single numerical parameter, that defines the ground truth protocol. The variability does not affect the outcome of the approach and rather captures the reality of variance in fiber behavior. For a visual overview of the stimulation protocol, schematic representations of all recordings are provided in Supplementary Figure S1.

**Table 1.** Ground truth protocol characteristics across recordings. Summary of stimulation protocol parameters for each dataset, including the number of additional pulses, inter-stimulus frequencies, and the distribution of temporal distances between the last additional pulse and the nearest background stimulus.

| Dataset | Additional pulses per interval (range) | Inter-stimulus frequency of extra pulses (Hz) | Distance to next background stimulus (range, ms) |
| --- | --- | --- | --- |
| A1 | 3-9 | 3-5 | 1000-3190 |
| A2 | 3-9 | 4 | 1730-3260 |
| A3 | 3-7 | 5-10 | 2600-3700 |
| A4 | 3-7 | 2 | 500-2500 |
| A5 | 3 | 2-8 | 3000-3670 |
| A6 | 1-7 | 5-8 | 3100-3700 |

### Recording with pruritogen injection

To illustrate the practical applicability of our computational approach to spike detection and sorting, we included one representative recording in which electrical background stimulation was combined with intracutaneous injection of 50 µL bovine adrenal medulla peptide 8-22 (BAM 8-22, Cat. No. SML0729, Sigma, Taufkirchen, Germany), as previously explained in [8]. Unlike the ground truth datasets, this recording did not include labels for spikes elicited by additional activity, making it a realistic illustration of the challenges posed by the detection and sorting of chemically induced spikes.

### Ground truth labeling with Dapsys via manual spike sorting

Spike identification and sorting for background stimulation-evoked activity were performed using the integrated tracking algorithm in Dapsys [24], followed by human expert corrections. In the ground truth recordings, this approach enabled the detection and sorting of spikes elicited by both background and additional stimuli. In contrast, for the recording with chemical stimulation, only spikes time-locked to background stimuli could be reliably identified with Dapsys.

### A knowledge- and data-driven spike sorting pipeline

We developed a Snakemake-based [25] computational pipeline for microneurography recordings (see Figure 3), integrating knowledge-driven steps (for example, background spike alignment and constraining the detection space) with data-driven machine learning for spike detection and classification.

**Figure 3.**
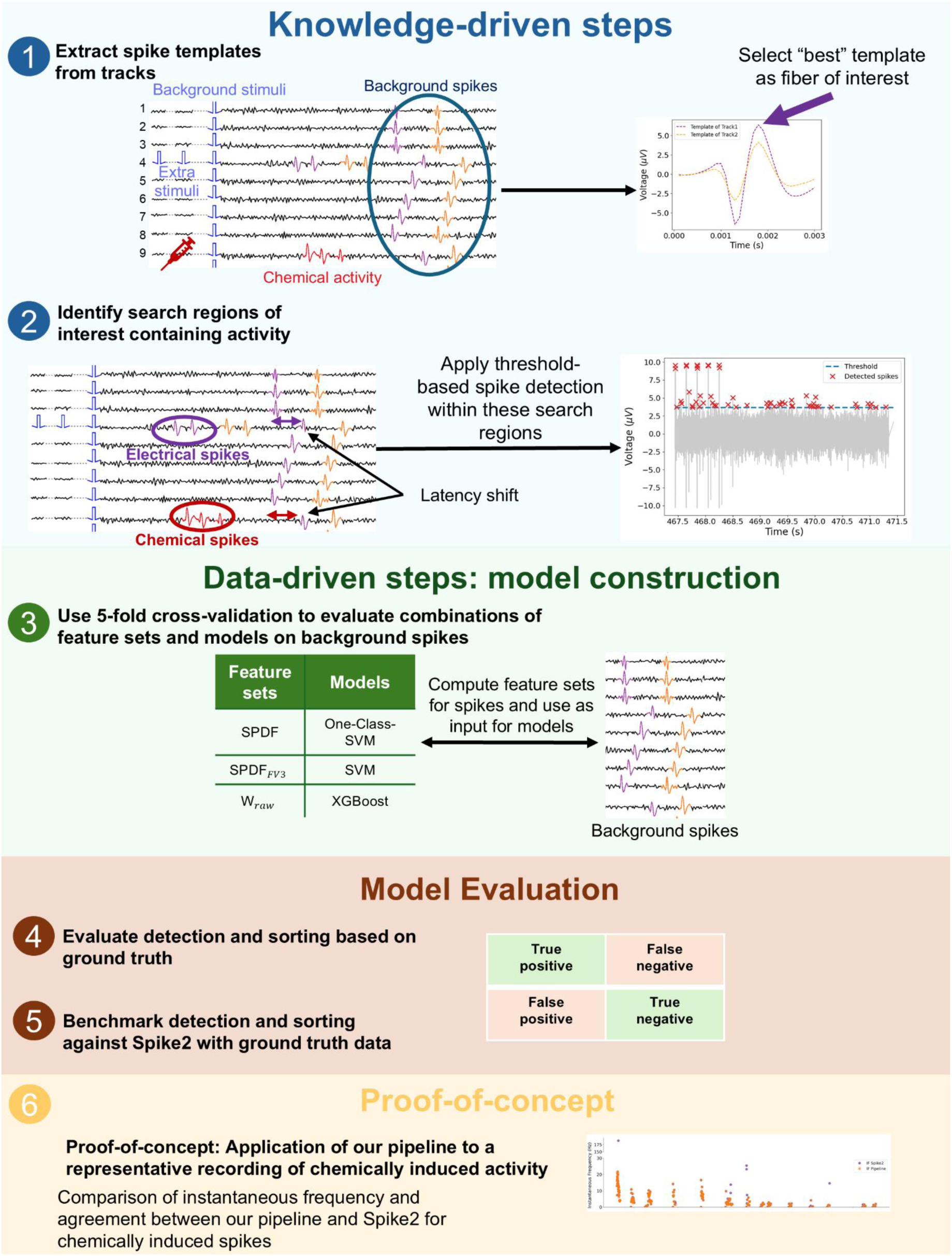
Overview of a hybrid spike detection and evaluation pipeline combining knowledge- and data-driven steps. (1) Spike templates are extracted from background spikes (blue circle) using low-frequency stimulation to identify the track of interest for further analysis. (2) Search spaces containing activity are identified, and spikes are detected using threshold-based methods. (3) Feature sets are computed from background spikes and used as input to different classification models (One-class SVM, SVM, XGBoost). Model performance is evaluated through 5-fold cross-validation to determine the optimal model-feature set combination for the background spikes. (4) Evaluate detection and sorting based on ground truth. First, spike detection is assessed by comparing detected spikes to known ground truth spikes, classifying outcomes as true positives, false positives, false negatives, or true negatives. Next, sorting performance is evaluated by verifying whether detected spikes are correctly assigned to their originating fibers. (5) Benchmark detection and sorting against Spike2 with ground truth data. The pipeline’s detection and sorting performance is directly compared to Spike2 by applying both methods independently to the same recording segments. (6) Proof-of-concept assessment of chemically evoked spikes. For the recording with pruritogen injection, we compute the instantaneous frequency, spike rate in 4-second windows, and latency of chemically evoked spikes sorted by both the pipeline and Spike2. As no ground truth is available for these spikes, this analysis serves as proof-of-concept, comparing both methods for the plausibility and physiological consistency of the detected activity and the agreement between both sorting approaches.

### Data pre-processing

All recordings were read in and parsed into structured pandas DataFrames (v2.2.3) containing the raw signal, spike labels, and timestamps, and simulation employing our custom Python package PyDapsys [26].

Spike waveforms were extracted from the raw signal with a fixed window of 30 data points, corresponding to a 3 ms interval at a sampling frequency of 10,000 Hz. Waveform alignment was performed according to the method described by Caro-Martín et al. [27], where each spike is temporally centered based on the time point corresponding to the largest downward slope (i.e., the negative peak in the first derivative of the signal). This alignment step ensures that extracted spike features are computed consistently across spikes.

### Template-based identification of optimal units

For each track, a template was computed by averaging all spikes aligned on that track. This template serves both as a visual representation of spike morphology and as a quantitative measure of spike amplitude, aiding in the identification of high-amplitude tracks where thresholding-based detection is more reliable. The track with the highest amplitude template was selected as the track of interest for subsequent spike sorting.

To assess the general quality, we computed the signal-to-noise ratio (SNR) for the track of interest. A noise segment of 40 ms was extracted from the raw signal preceding the first evoked spike to ensure the absence of fiber activity. The SNR was then calculated as the ratio of the spike template amplitude to the noise standard deviation, estimated using the median absolute deviation (MAD), following the implementation in SpikeInterface [16].

Spikes were further categorized by their consecutive temporal differences. Background-evoked spikes were defined as those occurring at least 3.8 seconds apart, consistent with the 4-second interval of the stimuli. In contrast, spikes with shorter inter-spike intervals were classified as responses to additional stimulation. Notably, background-evoked spikes are consistently available in recordings obtained via the marking method, whereas spikes evoked by additional stimuli are exclusive to datasets acquired using the ground truth stimulation protocol.

Template distance analysis was performed following our previously described method for estimating template separability and expected sortability [19]. We quantified the similarity between spike templates across tracks. Pairwise template distances were computed for all background-evoked templates using standard error-based metrics, including mean squared error (MSE), root mean squared error (RMSE), and mean absolute error (MAE). The definitions of the RMSE, MSE, and MAE metrics are provided in the Supplementary Methods. For each recording, the minimum distance across all template pairs was retained as an indicator of the most overlapping templates. Low template distance values indicate high similarity between spike waveforms and reduced separability and were therefore interpreted as reflecting lower expected sortability. These template-based distance measures were used as a pre-sorting quality indicator and later related to the measured F1-scores of the sorting result.

### Waveform drift test

For recordings exhibiting long recording durations, waveform drift can influence spike classification. We evaluated template stability over time for the selected track of interest evoked by background spikes. Spike templates were computed separately for the beginning, middle, and end of each recording and compared to assess potential changes in the spike shape. The track was divided into three subsets and all spikes within each segment were averaged to generate templates representing the beginning, middle, and end of the recording. These comparisons serve as a quality-control measure to confirm that the templates remained stable during the recording.

### Spike detection with constrained search space

Spike detection thresholds for the fiber of interest were set manually based on visual inspection of the peak amplitudes in the corresponding templates of background-evoked spikes. To establish these thresholds, the peak amplitudes were first sorted in ascending order to facilitate the identification of the minimum threshold at which spikes occur, and the resulting distribution was examined for variance and outliers. In recordings exhibiting high variance, we manually selected the threshold, otherwise the smallest amplitude value was selected. The final thresholds used are reported in Table 3. We intentionally decided not to use fully automated thresholding algorithms, as such methods are often sensitive to fluctuations and noise. Manual threshold selection allowed careful tailoring to the specific signal characteristics of each recording, thereby reducing the rate of false-positive spike detections.

**Table 2.**
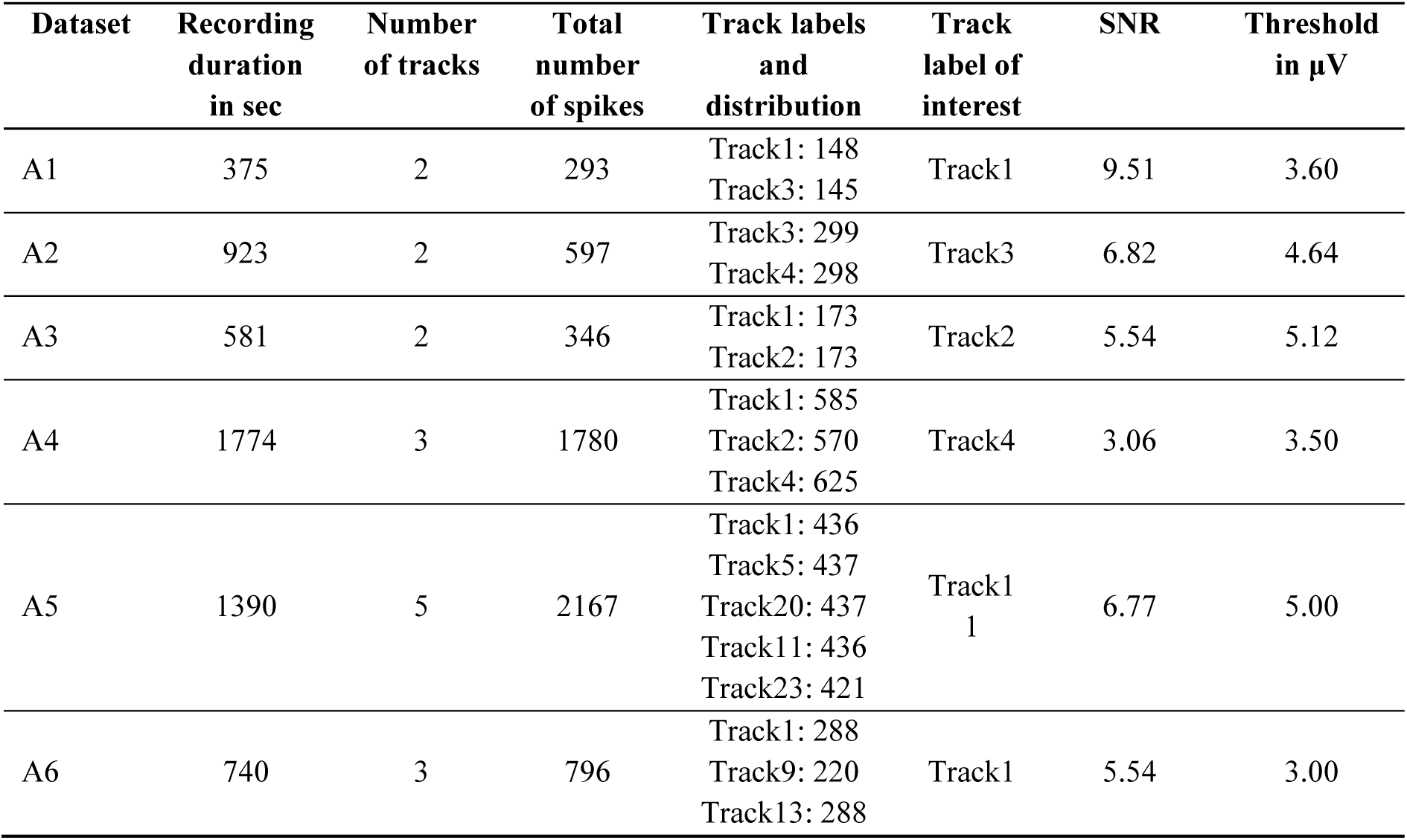
Data overview. The table summarizes the key properties of each dataset, including the total recording duration, the number of tracks (i.e., distinct recorded nerve fibers), and the total number of extracted spikes. For each track, individual spike counts are listed. Additionally, we specify the track selected for detailed analysis and the signal-to-noise ratio (SNR) computed for the track of interest.

**Table 3.**
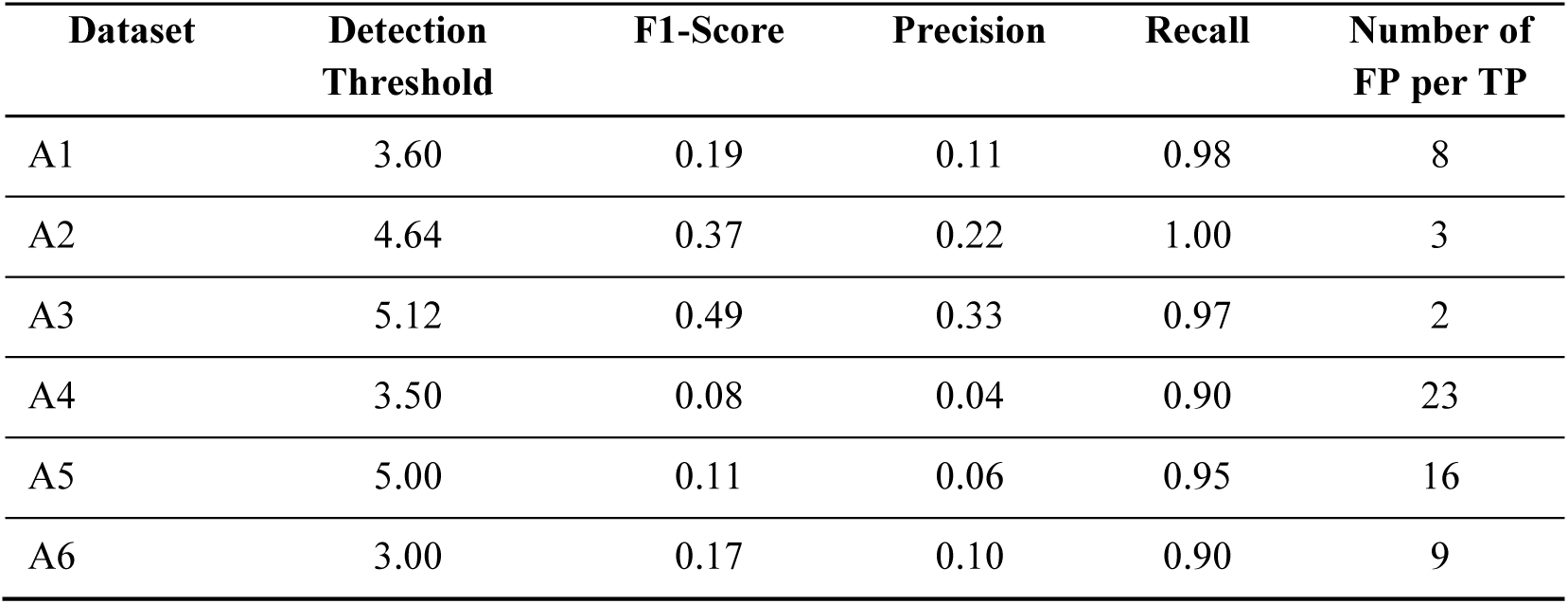
Spike detection performance via thresholding. For each recording, the detection threshold for the Dapsys recordings, F1-scores, precision, and recall are reported, along with the number of FP/TP. We set the thresholds based on the amplitudes of the spike templates and optimized the balance between reducing the number of false negatives and false positives.

To constrain the search space for threshold-based spike detection, we first calculated the spike latencies relative to the background stimuli for the track of interest. From the distribution of these latencies, we identified the minimal latency deviation, when no additional stimulation was applied, which was 0.9 ms. Any latency shift exceeding this value was then used to mark potential time windows within the 4-second signal intervals. These shifts indicate the presence of spikes evoked by additional stimuli and therefore constrain the search space for spike detection.

Spike detection was then performed using the *find_peaks* function from SciPy (v1.15.2) [28], applying the manually determined thresholds. To exclude stimulation artifacts, we excluded any detected peaks occurring within 100 ms of the stimulation onset.

### Spike classification using machine learning

To assign detected spikes to their corresponding track, we tested three machine learning classifiers: a one-class support vector machine (One-class SVM) and a standard support vector classifier (SVM), implemented in scikit-learn (v1.6.1) [29], and eXtreme Gradient Boosting classifier (XGBoost, v3.0.2) [30]. Feature vectors were extracted from the 30 datapoints describing the raw waveform. Three feature sets were computed: SPDF, a 23-dim feature vector, adjusted and implemented from the SS-SPDF features presented by Caro-Martín et al. [27], SPDF_FV3_, a reduced three-dimensional subset of SPDF, and W_raw_, the raw waveform feature vector, the 30 datapoints extracted from the raw signal. Detailed definitions of these feature sets are described here [19].

For the SVM and XGBoost classifiers, models were trained on background-evoked spikes from all available tracks in a recording, enabling multi-class discrimination across fibers. The One-class SVM model was trained using only background spikes from the track of interest, aiming to identify whether newly detected spikes match the known distribution of that track. Hyperparameter optimization for all models was performed using the Optuna framework (v 4.3.0) [31], applying 5-fold cross-validation on the background spikes. Maximizing the mean F1-score across folds was used as the optimization objective.

### Validation and evaluation methods

Detection and sorting performance were evaluated independently. For both, performance metrics included true positives (TP), true negatives (TN), false positives (FP), and false negatives (FN) rates, from which F1-score, precision, and recall were derived.

In the case of detection, we additionally report the false positive per true positive (FP/TP) rate as an additional indicator for reliability. Detection performance was assessed by comparing the timestamps of detected spikes against ground truth spike annotations using a temporal tolerance window of 2 ms.

For sorting, the final models were retrained on all background spikes using the previously selected optimal hyperparameters. Each model was then applied to the set of detected spikes, after extracting each of the three feature sets from those spikes to serve as input. Classifier performance was assessed with spikes evoked by ground truth.

### Statistical Analysis

Data was collected from multiple datasets and different performance metrics (F1-score, precision, recall, TP, FP, TN, and FN) were calculated. The metrics F1-score, precision, and recall are bounded in [0, 1]. As such proportions typically deviate from normality, we considered models suitable for bounded data. The metrics TP, FP, TN, and FN are count data and we therefore applied separate modeling strategies. All statistical analysis were done in R.

### Data Inspection and Preprocessing

We first conducted exploratory analyses (histograms, boxplots) to examine the overall distribution of each metric and identify any potential outliers or anomalies. The bounded metrics often approached boundary values (0 or 1). To avoid issues with values exactly at the boundary, each 0 or 1 was shifted slightly (ε = 0.0001) into the interval (0, 1).

### Comparing Statistical Models

For the bounded metrics, we compared two different statistical models: a Linear mixed model with logit transformation and a Beta mixed model, explicitly suitable for [0,1] data, fitted with a logit link. The model formula was expressed by OutcomeMetric ∼ Model ∗ FeatureSet + (1 | Dataset). For the count data, we compared a Poisson mixed model with a log link with a negative Binomial model. To test for overdispersion, we calculated the Pearson residuals and their dispersion ratio (sum of squared Pearson residuals divided by the residual degrees of freedom). The model formula was expressed by OutcomeMetric ∼ Model ∗ FeatureSet + offset(log(RecordingDuration)) + (1 | Dataset).

To compare the different models for the bounded as well as for the count data, we compared the information criteria (AIC/BIC) and conducted an analysis of the plots of the residuals compared to the fitted values.

### Baseline comparison with Spike2

To create a baseline for our algorithm, spike detection and sorting were also performed with Spike2 (v10.08), which implements a template-matching approach. The "New WaveMark" function was used with default parameters to define a reference spike, based on negative peak thresholds. This reference spike corresponds to a response evoked by a background stimulus and was temporally identified by the latency obtained from Dapsys, ensuring accurate association with a selected track. The identified waveform served as the initial template for spike sorting. Spike2 applied a template to detect and sort all subsequent spikes. The data was exported as spreadsheets and further processed in pandas DataFrames.

To remove stimulation artifacts from the Spike2 result, spikes occurring within a 10 ms window following each stimulus pulse (stimulus onset + 10 ms) were excluded.

Simultaneously recorded datasets from Dapsys were used to obtain ground truth labels. A key challenge was the temporal misalignment between the two systems: Dapsys recording sessions were segmented across multiple files, while Spike2 generated a single recording file.

A dynamic offset correction was applied to align Spike2 spike timestamps with their corresponding Dapsys timestamps. This offset was computed individually for each stimulus, accounting for time drift between systems.

After alignment, spike sorting performance was assessed by comparing Spike2-detected spikes against Dapsys ground truth labels. A Spike2 spike was considered a true positive if it occurred within ±3 ms of a Dapsys spike. This tolerance window corrected inherited differences in timestamp definitions: Spike2 timestamps correspond to the spike onset, whereas Dapsys timestamps approximate the midpoint of the spike. This evaluation enabled the assessment of Spike2’s template-based sorting performance relative to the manually labeled Dapsys result.

### Proof-of-concept comparison of chemically evoked spikes in the absence of direct ground truth

To illustrate the application of the computational spike sorting pipeline under experimental conditions without direct ground truth, we performed a proof-of-concept comparison with Spike2 for a representative recording with pruritogen injection. Spikes assigned to the fiber of interest by both methods were compared to assess the degree of agreement between the sorting approaches. The instantaneous firing frequency (IF) was calculated as the inverse of the interspike interval between consecutive spikes and the spike rate was additionally quantified in 4-second time bins. These measures were used as an indirect measure of fiber activity evoked by chemical stimulation, enabling them to examine the consistency between the two sorting approaches. To further characterize the spike sorting quality, spike latency relative to the background stimuli was visualized as a physiological marker of fiber activation. Latency shifts were plotted alongside instantaneous frequency and spike rate to detect inconsistencies and assess the plausibility of detected bursts in both methods.

### Software Accessibility

The complete spike sorting pipeline, including data preprocessing, feature extraction, and classification, is openly available at https://github.com/Digital-C-Fiber/SpikeSortingForSpikingPatterns. The repository includes source code, documentation, and an example recording.

## Results

### Data characteristics

In this work, we systematically evaluated spike sorting performance across six microneurography recordings, including challenging scenarios with low signal-to-noise ratios (SNR) and a high number of nerve fibers. The recording durations range from 375 to 1,774 seconds with SNR values between 5.13 and 9.20, and the number of tracks varies from 2 to 5. Table 2 provides a comprehensive summary of the recording characteristics. By selecting these diverse recordings, our analysis provides a robust test of spike sorting methodologies under varied and demanding conditions. To facilitate the identification of a track of interest, we computed a template representing the average of all tracked spikes for each recording.

Figure 4a displays these templates. Additionally, to provide an overall impression of the signal quality and recording conditions, we computed the SNR for each dataset (see Table 2). A comparison of the computed template with a short segment of background noise is illustrated in Figure 4b, highlighting the differences between signal and noise characteristics. This visualization offers an immediate qualitative impression of the recording challenges encountered in each dataset. To assess waveform drift over time, spike templates were compared across the beginning, middle, and end of each recording. Minor changes were observed in a few recordings, for example, in A2, the maximum template amplitude increased slightly from ∼7 μV at the beginning to ∼8 μV at the end, and in A5, a slight tilt appeared in the middle and end templates. These deviations were very small and are fully captured in the sorting process by background-evoked spikes used as training data. Overall, no substantial waveform drift was detected in any of the six datasets (see Supplementary Figure S2), indicating that spike waveforms remained stable throughout the recordings.

**Figure 4.**
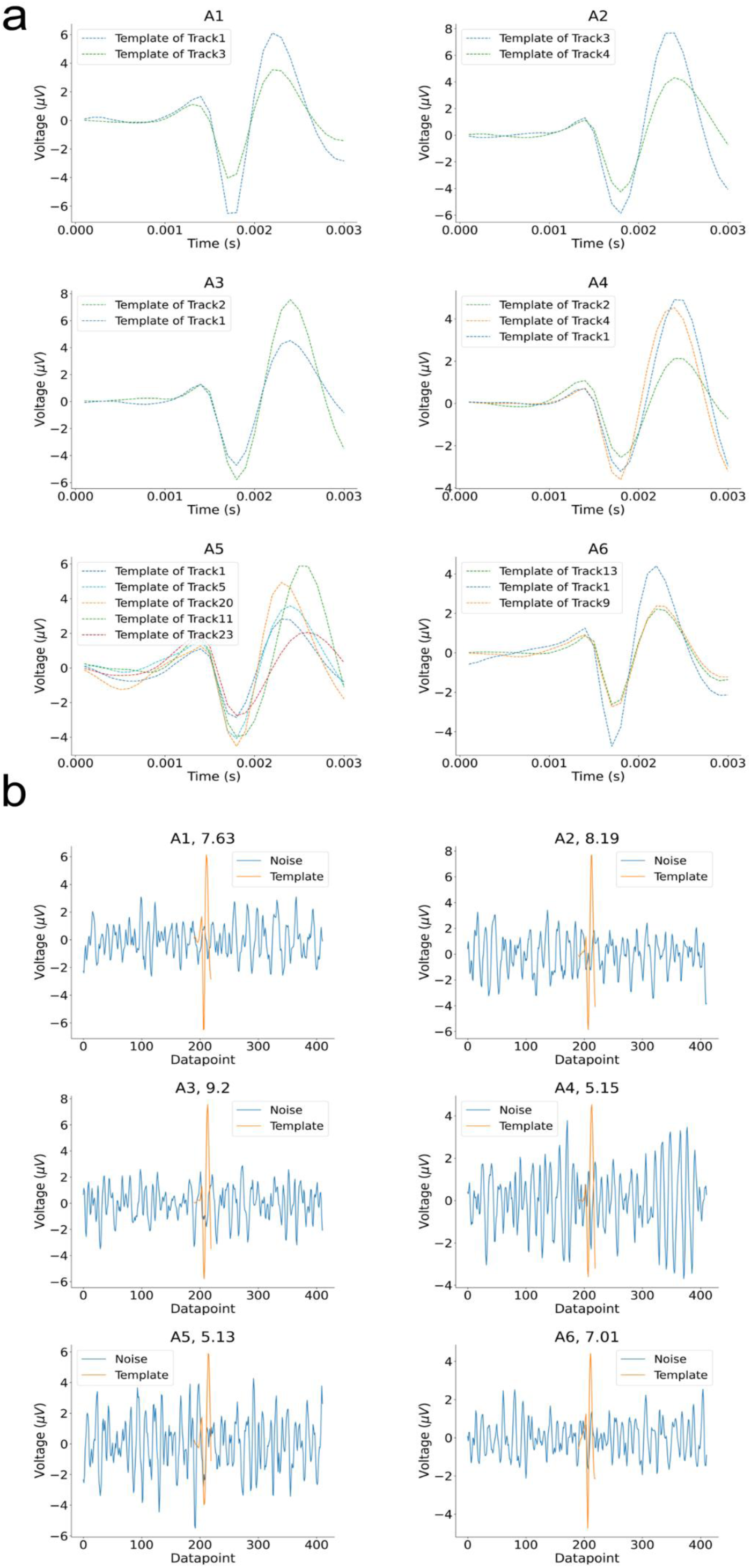
(a) Average spike templates for each track in all recordings. For each dataset, we extracted spike waveforms from each track and aligned them according to the time point corresponding to the largest downward slope (time point of the maximum negative peak in the first derivative of the signal as the temporal midpoint) [22]. We averaged all spikes to generate the templates. Each subplot shows the templates for all tracks within a single recording. These templates reflect the characteristic spike shapes, enable the visual comparison of waveform morphology, and support the selection of the track of interest. (b) Signal-to-noise (SNR) visualization for each recording and track of interest. For each recording, the track containing the spikes with the largest amplitudes was used to calculate the SNR. We calculated the SNR by dividing the peak amplitude of the template by the standard deviation of a noise segment extracted from the raw recording. The noise standard deviation was estimated using the median absolute deviation (MAD). The computed SNR for each dataset is indicated next to its name in the figure title.

### A computational pipeline for adaptive spike detection and sorting

#### Constraining the search space with latency information

To evaluate the detection rate of our pipeline, we employed a thresholding approach to identify spikes in the recordings. The detection performance was analyzed for each individual recording. We constrained the search space, focusing on segments preceding a latency jump to improve detection results.

The absolute numbers of detected spikes are visualized in Figure 5. The false positive rate per true positive (FP/TP) varies significantly across datasets. A4 and A5 exhibit the highest values for FP per TP with 20 and 14, respectively. Our approach demonstrated a high recall rate, reflecting the proportion of true spikes correctly detected, suggesting that by separating the spike detection and sorting steps, we can first reliably detect spikes before proceeding to classification.

**Figure 5.**
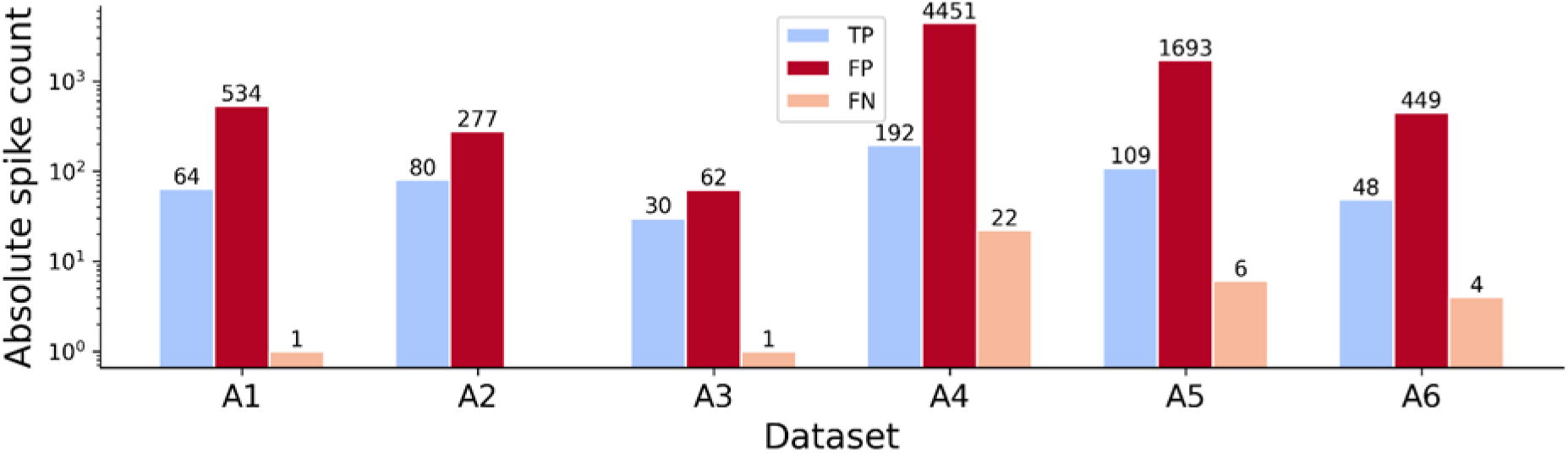
Spike detection performance using thresholding and peak detection. TP, FP, and TN are visualized on a log scale for the detected spikes in the constrained search space. All recordings have higher false positive values compared to the TP values. A4 exhibits the most false positives and 22 spikes could not be detected with the set threshold.

However, in certain cases, increasing the detection threshold proved beneficial and slightly led to a reduction in false positives, at the cost of tolerating a few more false negatives, for example, in the case of A4. This balance is crucial for optimal spike sorting performance. All detection thresholds and computed metrics are listed in Table 3. All metrics are collected in Supplementary Table S1.

#### Individual recordings require individual combinations of feature sets and classifying algorithms

After the initial detection step, we employed different input feature sets (SPDF, SPDF_FV3_, and W_raw_) and classification models (One-class SVM, SVM, XGBoost) for spike sorting. A comprehensive comparison of all tested approaches is presented in Figure 6a, which shows the values of F1-score, TP, FP, TN, and FN for each recording. The rectangle highlights the best-performing method for each dataset. The results clearly demonstrate that no single sorting approach yields universally optimal performance across all recordings. Instead, each dataset exhibits its own optimal configuration, underscoring the variability in recording conditions and variance for microneurography. Among the datasets, A1 consistently produced the highest overall performances across methods, while A4 presented the most challenging conditions, with generally lower F1-scores and more frequent misclassifications.

**Figure 6.**
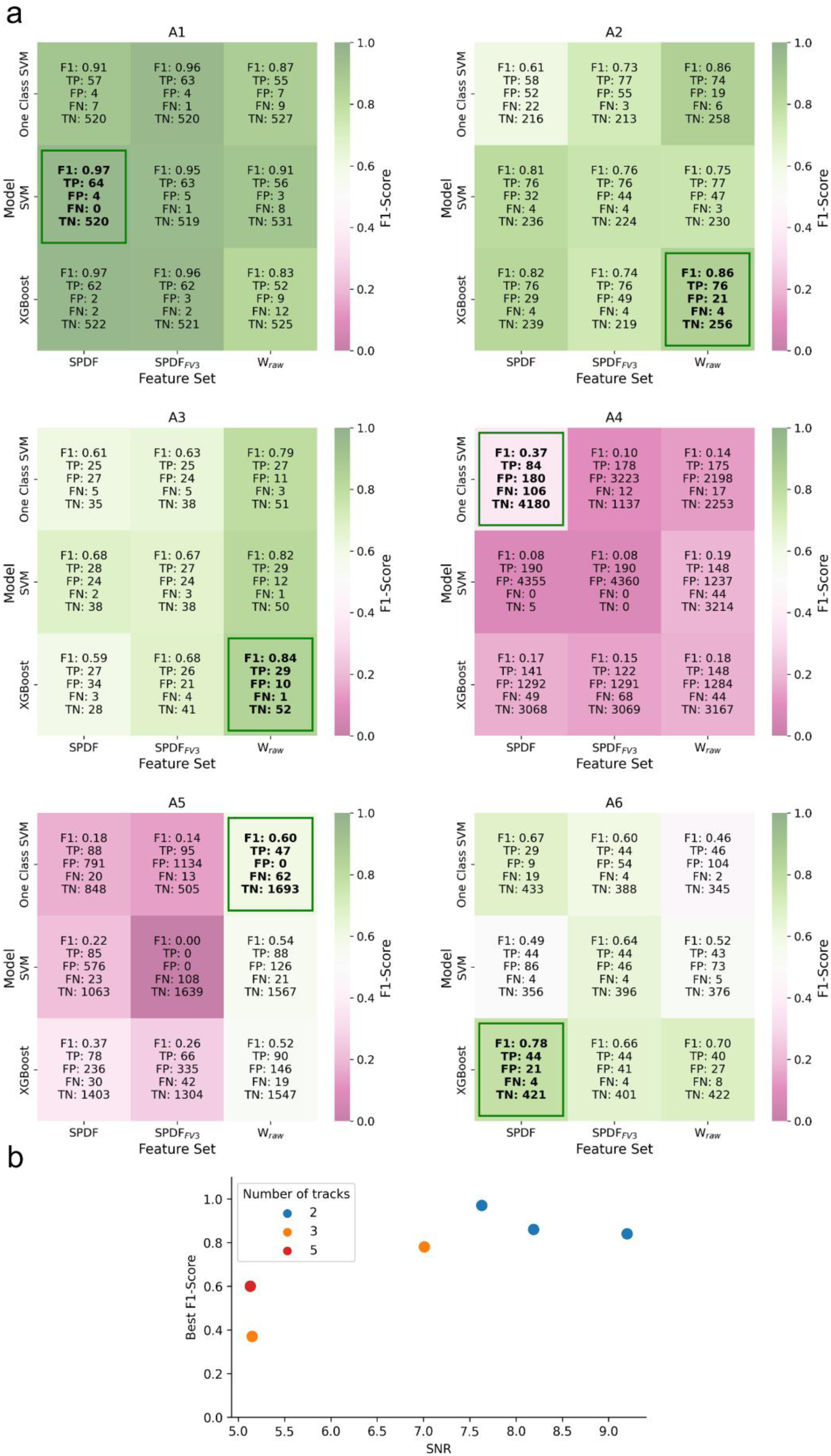
**(a)** Heatmaps of spike sorting performance across feature sets, classification models, and evaluation metrics. Each cell is color-coded according to the F1-score with magenta indicating low performance and green indicating high performance. The heatmaps display results for all tested combinations of three feature sets and classification models, evaluated using multiple metrics (F1-score, TP, FP, FN, and TN). The green rectangle highlights the best-performing configurations based on the F1-score for each recording. **(b)** Relationship between SNR and best F1-score for recordings with a varying number of tracks. Each dot represents a recording colored by the number of simultaneously recorded tracks (2, 3, or 5). As expected, lower SNR values correspond to reduced F1-scores. Notably, the F1-score plateaus beyond an SNR of approximately 7.5, indicating no further improvement in performance.

Similar to our findings in the detection phase, the sorting results show strong dataset dependency, with performance varying considerably across different recording conditions and microneurography signal characteristics. To investigate how recording properties influence sorting performance, Figure 6b presents the relationship between SNR and the best F1-score. As expected, recordings with lower SNR exhibit lower F1-scores, demonstrating a clear dependency between noise levels and classification performance. However, beyond an SNR of approximately 7.5, the F1-score no longer improves.

Recordings with fewer fibers generally yield higher F1-scores with all recordings containing a small number of fibers achieving scores above 0.8. Interestingly, the dataset with the highest number of fibers (A5) does not produce the worst result.

These findings suggest that improving the signal quality and reducing the number of active fibers in an active recording session are crucial for good sorting results, which should be considered during the experiments. To further characterize the variability in sorting performance and SNR across models and feature sets, we computed the 95% confidence intervals for each dataset using SciPy’s bootstrap resampling across all feature set-model combinations (see Figure 7). This analysis shows the stability of different approaches within each dataset. Consistent with the results observed above, the two datasets with the lowest SNR (A4 and A5, SNR ≈ 5) exhibit the weakest and least reliable performance. For A5, the confidence interval is the widest among all datasets, indicating uncertainty and strong dependence on the specific model-feature set combination. In contrast, A4 displays consistently low performance with a narrower interval, confirming that nearly all evaluated models fail on this dataset.

**Figure 7.**
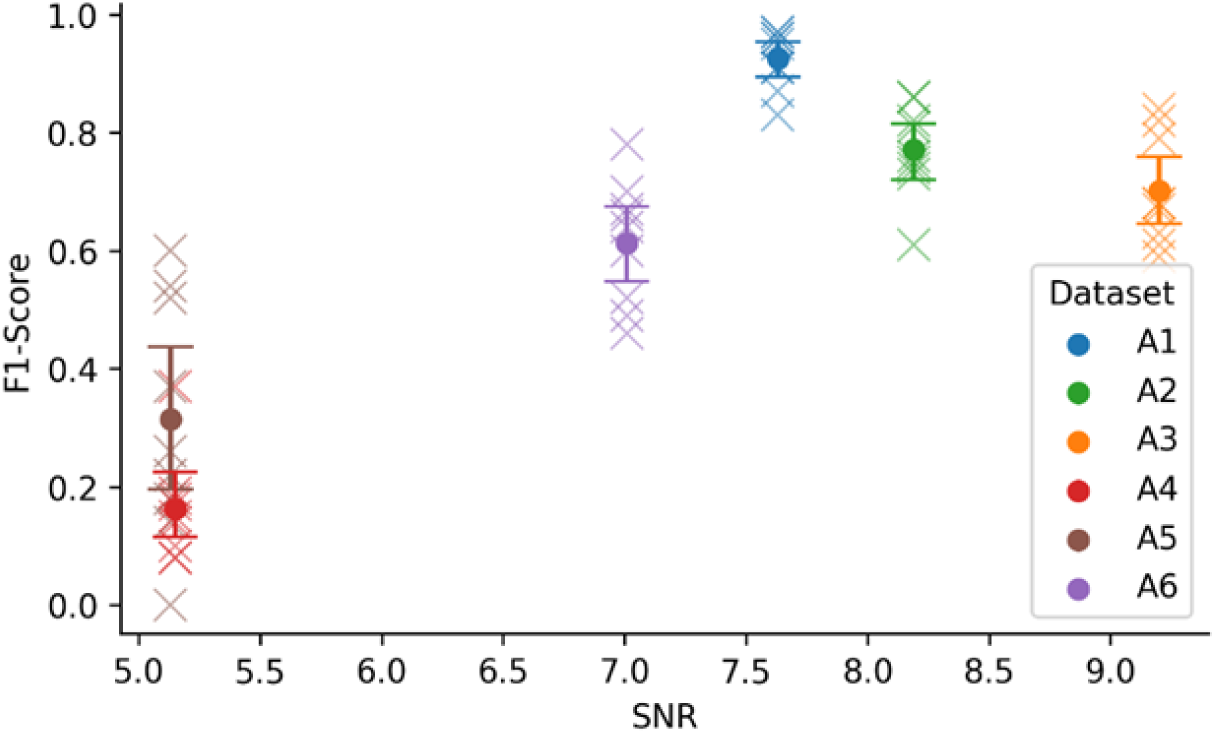
F1-scores and confidence intervals for all datasets. Each cross shows an F1-score obtained from one model-feature set combination. The error bars and the dot indicate the mean for the specific dataset and the ±95% confidence intervals computed by non-parametric bootstrap resampling across all combinations.

For the higher SNR datasets, the confidence intervals do not overlap with those of A4 and A5, indicating distinct performance distributions. A3, the recording with the highest SNR, does not achieve the highest mean F1-score. This demonstrates that while SNR is a helpful indicator, it should not be used in isolation to predict spike sorting quality. Nevertheless, all higher SNR datasets have relatively narrow confidence intervals, indicating stable performance across different combinations.

#### The combo XGBoost + W_raw_ achieves the highest median F1-score

To evaluate the performance across different combinations of feature sets and models, we visualized the F1-score using boxplots (see Figure 8a) and a scatter plot (see Figure 8b). All metrics are collected in Supplementary Table S2. In addition to precision and recall, we report the false discovery rate (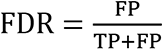) and the mean number of misassigned spikes per searched interval (FP/N_intervals_), computed as FP divided by the number of 4-second stimulation intervals for the best results of each recording (see Supplementary Table S3). For each feature set, XGBoost (orange) consistently achieved the highest median F1-score and demonstrated lower variability compared to SVM (green) and One-class SVM (blue). Among the feature sets, the raw waveform features (W_raw_) yielded slightly higher and more consistent F1-scores than both the full SPDF set and its reduced subset SPDF_FV3_.

**Figure 8.**
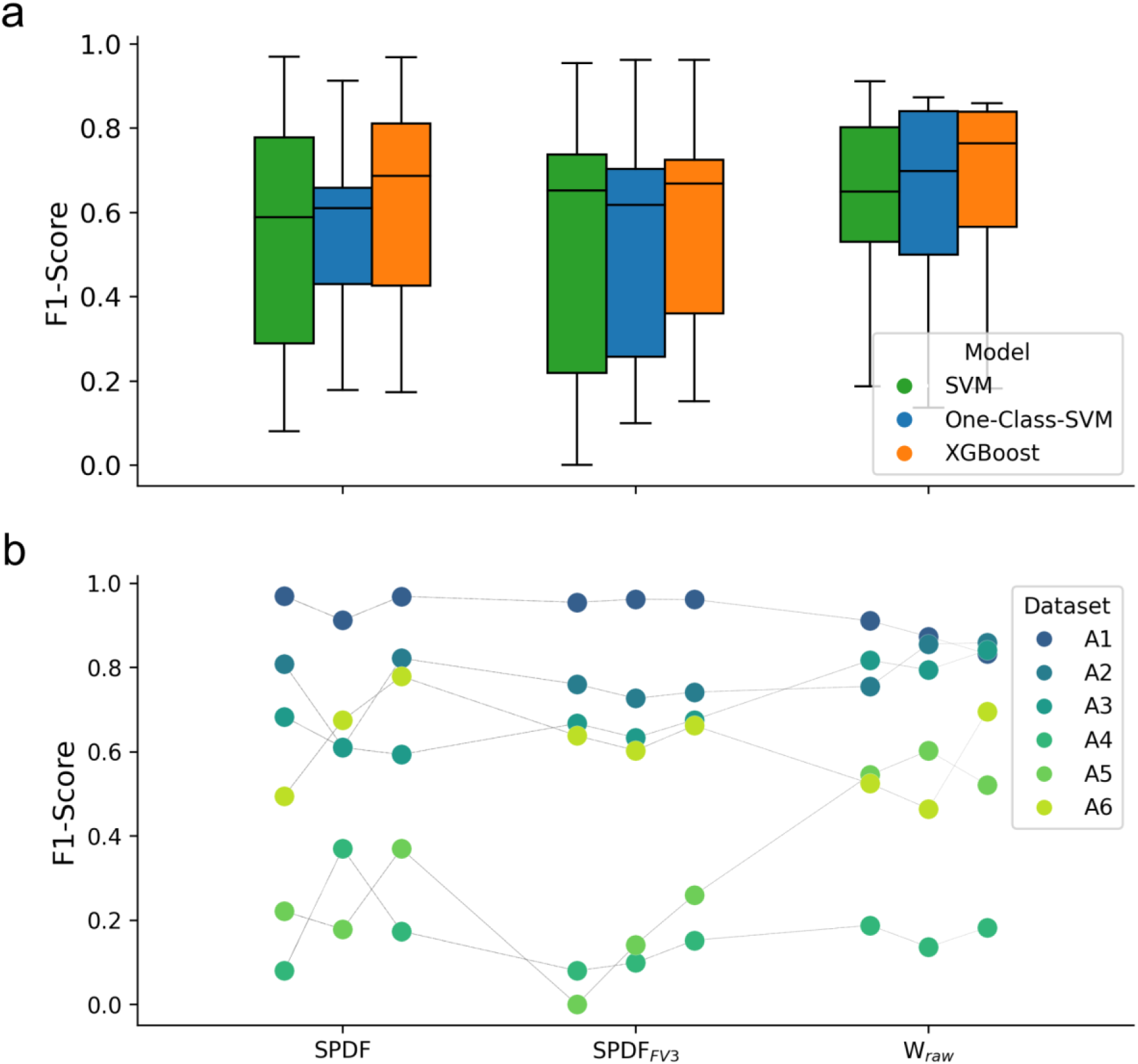
(a) Boxplots of F1-scores are shown for three feature sets (SPDF, SPDF_FV3_, and W_raw_), evaluated across three classification models: One-class SVM, standard SVM, and XGBoost. Each boxplot summarizes the distribution of F1-scores for all six recordings with the horizontal line indicating the median. (b) Dataset-specific trends are illustrated, showing F1-scores for each dataset (A1-A6) across the combinations. The grey lines connect scores for each dataset to illustrate relative changes in performance. While some recordings, such as A1, show good performance for all combinations, others show greater variability depending on the chosen feature set and model (for example, A5).

For the statistical analysis, the outcome measurements were divided into two categories: bounded metrics (F1-score, precision, and recall) and count metrics (TP, FP, TN, and FN). An initial assessment showed that none of the metrics followed a normal distribution.

Consequently, we fit different mixed-effects models appropriate to each metric type, as described in the Methods section. Model comparisons via information criteria (AIC/BIC) indicated that the Beta mixed model consistently provided the best fit for the bounded metrics, while the negative Binomial model exhibited the lowest AIC/BIC values and favorable dispersion ratios for count metrics. In both final models, plots of residuals versus fitted values revealed a random scatter around zero, suggesting no discernible patterns and confirming that variability in the data was adequately captured.

We applied the mixed-effects models to investigate the influence of the classification models, feature sets, and their interaction on various performance metrics statistically (see Table 4).

**Table 4.**
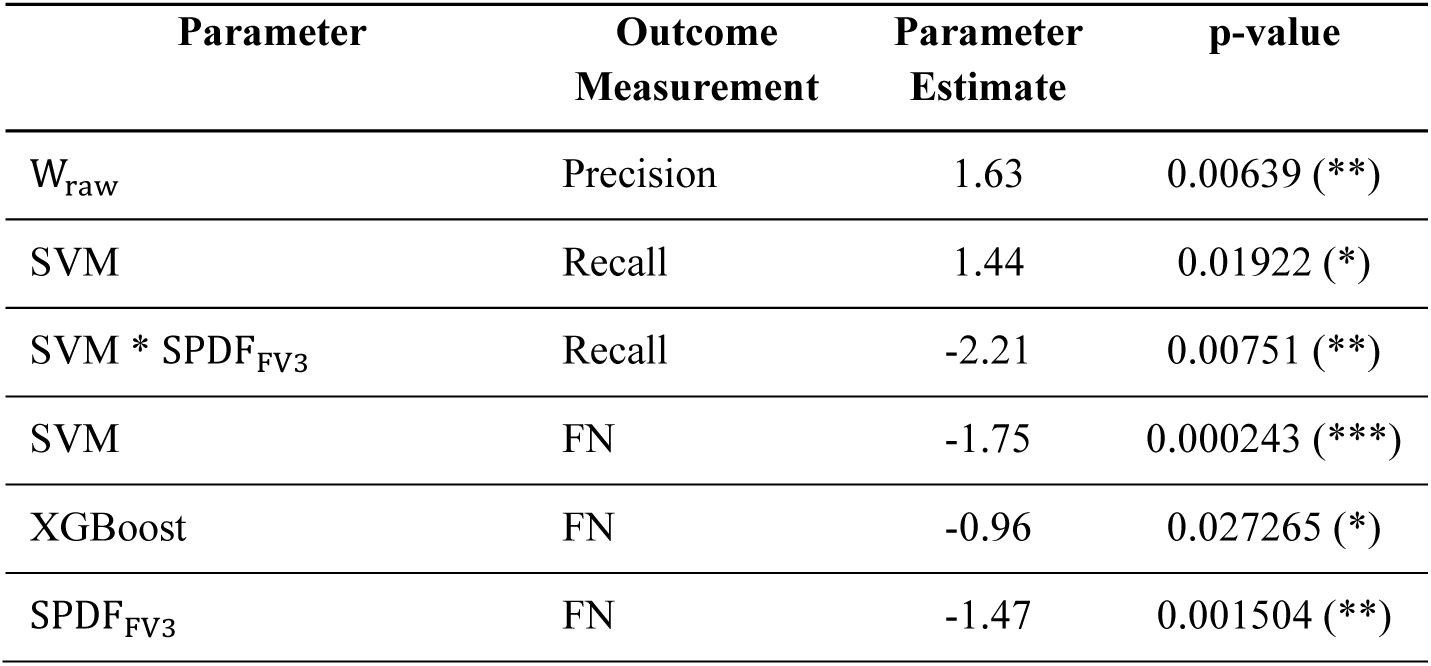
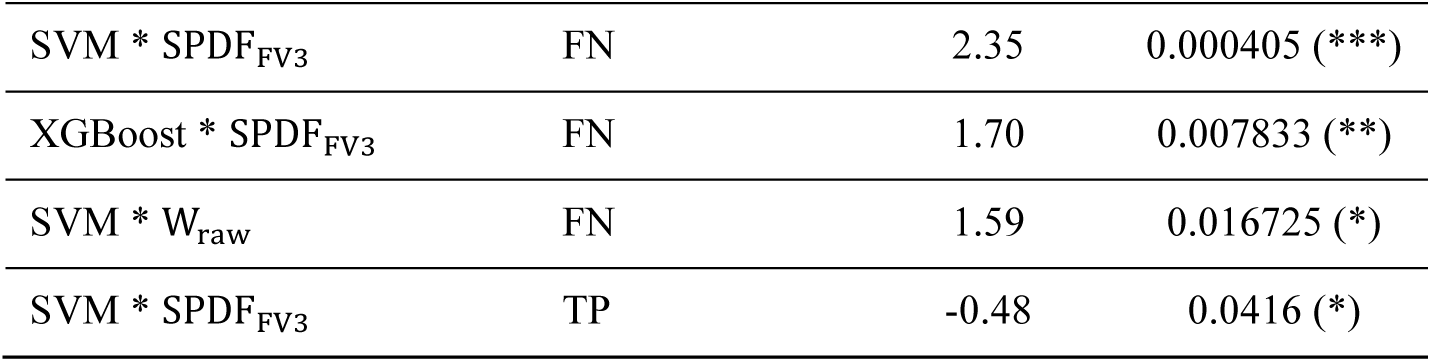
Statistically significant parameter estimates (p < 0.05) from the Beta and negative Binomial mixed models across the eight-outcome metrics (F1-score, precision, recall, FN, FP, TN, and TP). Estimates for F1-score, precision, and recall are presented on the logit scale. Estimates for FN, FP, TN, and TP are presented on the log scale. Non-significant terms are omitted for clarity. Significance levels are denoted by asterisks (*, **, ***).

The One-class SVM model and the SPDF feature set were used as a baseline and compared to the other models and feature sets, as well as their interactions. For the F1-score, no main effects or interactions reached statistical significance, suggesting relatively consistent performance across conditions. In contrast, false negatives showed multiple significant effects: both SVM and XGBoost models, as well as the SPDF_FV3_ feature set, significantly reduced FN counts compared to the baseline. Even though the interaction terms SVM * SPDF_FV3_, XGBoost * SPDF_FV3_, and SVM * W_raw_ increased FN significantly, the total effect, calculated by summing the main effects and the interaction effects for each combination, remained negative (SVM and SPDF_FV3_= -0.87, XGBoost and SPDF_FV3_= -0.74, SVM and W_raw_ = -0.92). A similar effect was observed for true positives, where the interaction between SVM and SPDF_FV3_ significantly reduced TP counts, but the total effect remained positive (0.05), indicating a potential trade-off between capturing more positives and avoiding false alarms. Models for TN and FP did not yield statistically significant predictors. These findings highlight that while overall F1-scores remain stable, specific configurations influence false positive and false negative distributions differently.

#### Template similarity as a pre-sorting indicator

Template distance analysis revealed a strong relationship between waveform similarity and the best sorting performance. Recordings with highly similar templates showed the poorest sorting performance. A4 exhibits the lowest F1-score of 0.37 and simultaneously shows the smallest distance values across all three metrics (see Figure 9), consistent with the interpretation that high template similarity makes fibers difficult to separate. A5 shows the second-highest error yet achieves only the second-lowest F1-score of 0.60. This discrepancy can be attributed to the high number of active fibers in the recording (five), highlighting that not only template similarity but also the number of fibers critically influence the sorting success. Together, these findings support the use of template-based distance metrics, combined with the number of active fibers and SNR, as a practical pre-sorting quality index for judging whether a recording is suitable for spike sorting.

**Figure 9.**
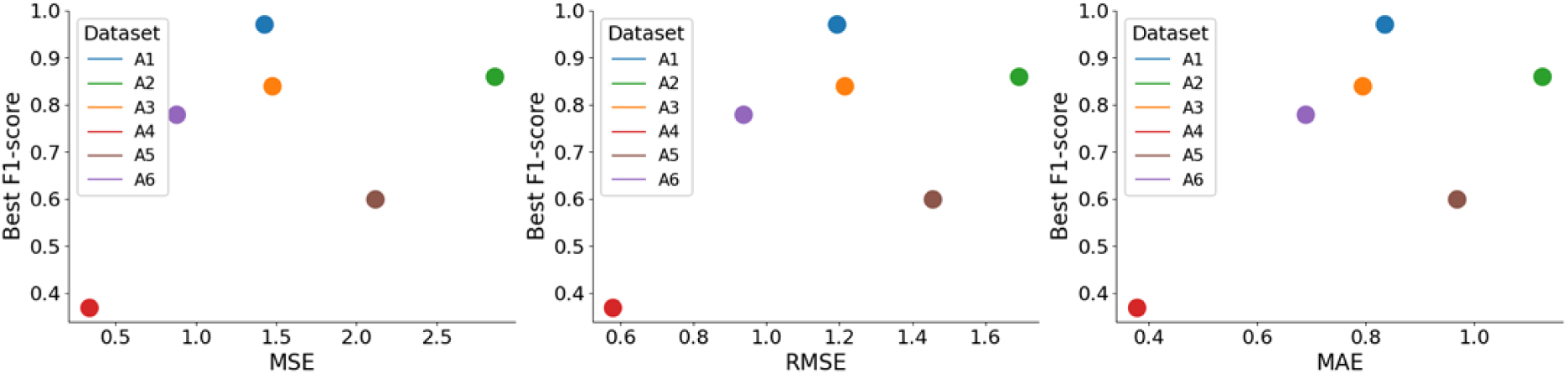
Relationship between template similarity and spike sorting performance. Scatter plots showing the relationship between template distance metrics computed between the two most similar fiber templates (minimal error) in each recording, and the corresponding best F1-score achieved by our spike sorting pipeline for all six datasets. Lower template distance values indicate higher similarity (greater overlap) between fiber templates and are associated with reduced sorting performance. In combination with fiber count, the relationship can provide a useful pre-sorting indicator of expected sorting reliability. (a) Mean square error (MSE), (b) Root mean squared error (RMSE), and (c) Mean absolute error (MAE).

#### Feature importance analysis

To identify which features contributed the most to the sorting result, we performed a feature importance analysis using two models: a random forest classifier, which captures nonlinear interactions between features, and an L1-regularized logistic regression model, which yields sparse linear coefficients. For each recording, model, and feature set, we computed the importance scores and extracted the ten highest-ranking features. The results are shown in Supplementary Figure S3.

Importantly, no single feature dominated across recordings, which is consistent with the results from our main analysis that each recording has its own optimal combination of model and feature set. The SPDF_FV3_ feature set produced compact groups of high-ranking features for some recordings, whereas the SPDF and raw waveform (W_raw_) feature sets showed broader and more scattered importance distributions. However, none of the feature sets exhibited a stable or interpretable feature ranking across all datasets.

These results demonstrate that feature importance is highly recording-dependent, mirroring the inherent heterogeneity of microneurography recordings. Rather than relying on a small set of universally informative features, classifier performance appears to arise from the interaction of multiple feature dimensions with different recordings, emphasizing different aspects of the signal.

#### Analyses using Spike2 are only reliable in high SNR recordings

As a baseline for comparison, we applied a standard spike sorting pipeline using Spike2 (v10.08), which processes the entire duration of each recording using thresholding and template matching. When applied to the full recordings (recording durations in Table 3), as intended by Spike2, most datasets exhibited a high number of FP (see Figure 10a, Spike2 (Full Recording)). The only exception was recording A1, where the number of TP exceeded false positives. This can be attributed to a higher SNR (7.63) and only two tracks. In contrast, recordings A4 and A5 showed the highest number of FP per TP (see Table 5), with A4 also exhibiting the highest false negative rate, approximately 25% of spikes were not detected and sorted. A5 contains activity from five tracks, which is a possible reason for the high number of TP but a relatively low number of FN. To avoid the influence of recording duration giving a false impression, we also report scaled standard metrics, such as F1-score, precision, and recall (see Table 5). While a promising recall remains around 0.9 for almost all recordings except A4, the poor overall performance becomes evident with a mean F1-score of 0.33 and a mean precision of 0.25 across all recordings. All metrics are collected in Supplementary Table S4.

**Figure 10.**
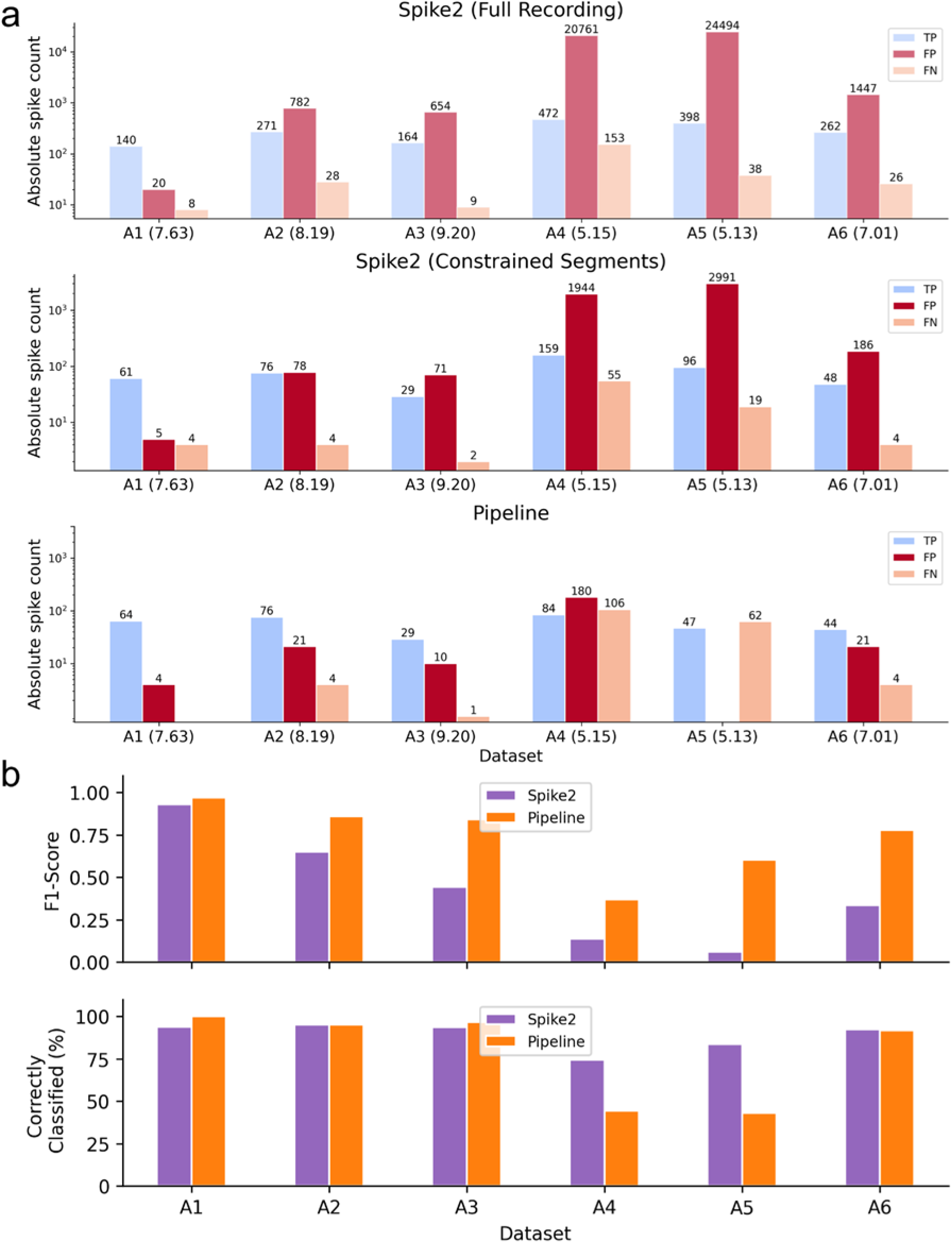
Spike detection and sorting performance comparison between Spike2 and our pipeline across datasets A1-A6. (a) Bar plots show absolute counts of TP, FP, and FN for each dataset on a log scale. The top panel shows results from Spike2 on the full recordings, the middle panel shows Spike2 results restricted to the same segments found via the pipeline, and the bottom panel shows results from our pipeline using the best-performing feature set-model combination (based on F1-score). The computed SNR for each dataset is shown in parentheses next to the dataset name. (b) Quantitative comparison of sorting performance between Spike2 and our pipeline. The top plot displays F1-scores, while the bottom plot shows the percentage of correctly classified spikes.

**Table 5.**
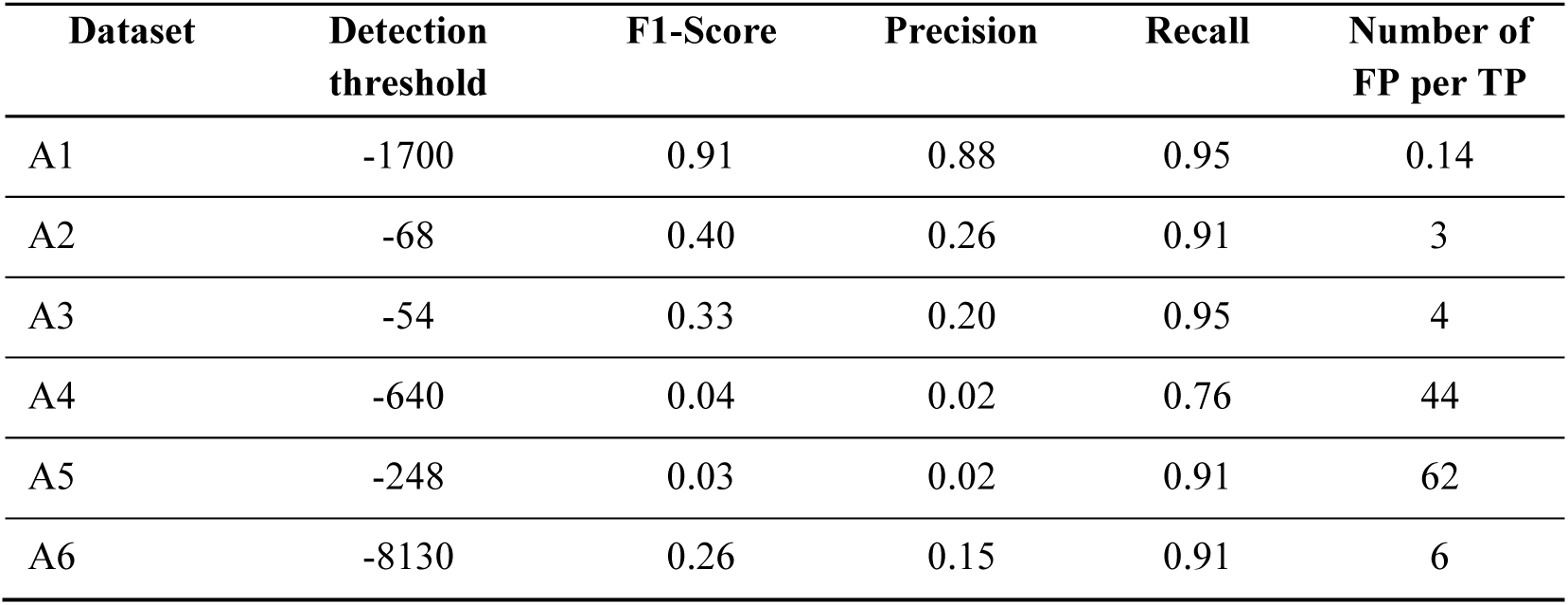
Spike sorting performance using Spike2. For each recording, the detection threshold used in Spike2 is reported along with the resulting F1-score, precision, and recall values. The last column indicates the number of FP/TP, providing a measure of spike sorting specificity. In well-sorted recordings, this value should ideally be below 0.5, indicating fewer than one FP for every two TPs.

Furthermore, a core challenge in the analysis of our results is the dynamics of spike waveforms during long recording sessions. Initially, a template was created for the spikes of the track of interest, while other spikes that fall within the threshold range but do not match the template trigger the creation of new templates. In theory, these additional templates could be ignored. However, as the experiment progresses, the waveform of the spike of interest changes. This shift means that some templates initially classified as incorrect may later contain spikes from the track of interest. Consequently, post-hoc deletion of these later templates would result in missing relevant spikes. Table 6 shows an example for A2.

**Table 6.**
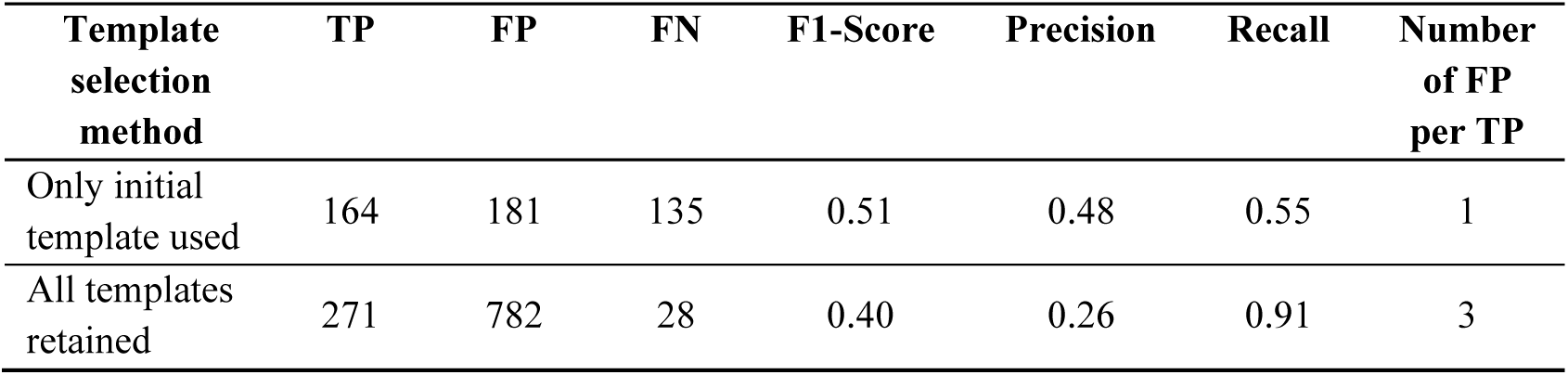
Impact of template selection strategy on performance in recording A2. The comparison of performances for two approaches is shown: using only the initial spike templates vs. retaining all templates generated during the analysis. We summarized TP, FP, FN, F1-scores, precision, and recall values for both strategies. Due to waveform drift over time, retaining all templates improves the spike detection and sorting that would otherwise be missed (recall improves to 0.91 from 0.55).

This issue becomes particularly evident now that we have access to ground truth data, allowing us to verify how template evolution affects classification. In real-life scenarios, without ground truth validation, we would have to accept the limitations of a single-template approach, which would lead to an increased number of FNs. Alternatively, accepting all detected templates would capture more TP but at the cost of potential misclassification. This trade-off highlights the challenges in spike sorting and the difficulty in determining the optimal balance between precision and recall.

Based on these results and the observed recall values, we decided to retain all templates. Without ground truth data, it is impossible to determine with certainty which spikes belong to the fiber class and which do not. Prioritizing high recall ensures that we capture as many true spikes as possible, even at the cost of more false positives. While false positives could possibly be removed later through additional filtering or manual inspection, missed spikes cannot be recovered, making this approach a more reliable choice for our analysis.

To enable a direct comparison with our own pipeline, we restricted the search space to the same intervals where we observed latency jumps (see Figure 10a, Spike2 (Constrained Segments) and Pipeline). All metrics are collected in Supplementary Table S5.

In recordings with high SNR (A1-A3, A6), our pipeline demonstrated a clear advantage over Spike2, correctly identifying and sorting the majority of spikes while generating fewer false positives. In contrast, Spike2 exhibited higher false positive rates. Under more challenging recording conditions, for example, A4 and A5, both approaches showed limitations: Spike2 continued to produce a large number of false positives, whereas our pipeline was more restrictive, which resulted in fewer incorrect classifications but missing a substantial number of spikes, reflected as an increase in false negatives.

To provide a final performance comparison, we evaluated both F1-score and the proportion of correctly classified spikes for each dataset (see Figure 10b). Our pipeline outperformed Spike2 in terms of F1-scores across all datasets, indicating a better balance between true positive and false positive classifications, though Spike2 yielded a higher percentage of correctly classified spikes in A4-A6. Notably, for dataset A1, both methods achieved similar performance with nearly equivalent F1-scores and high classification accuracy. Spike2 maintained a higher percentage of correctly classified spikes, likely due to its tendency to classify more events overall, including many false positives. This highlights a key difference: our pipeline establishes conservative but precise classification, whereas Spike2 prioritizes higher coverage at the expense of precision.

Using the described mixed-effect models from before, we compared each classification method against Spike2 (see Table 7). One-class SVM and XGBoost significantly outperformed the Spike2 baseline across multiple outcome metrics (F1-score, precision, and FP). The standard SVM showed a significant improvement only in terms of false positives (FP).

**Table 7.**
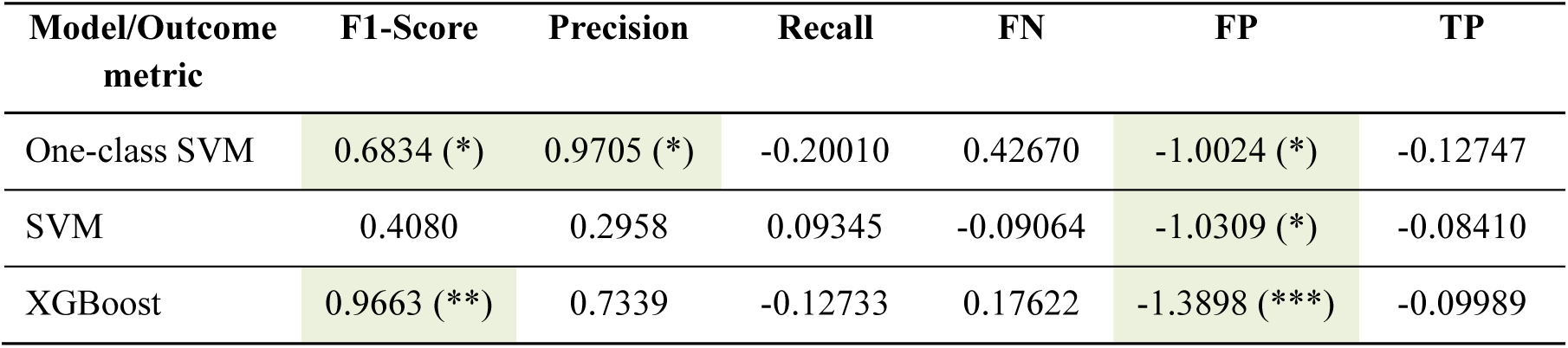
Parameter estimates (with significance levels) for each model (One-class SVM, standard SVM, and XGBoost) compared to the Spike2 baseline across six outcome metrics (F1-score, precision, recall, FN, FP, and TP). Estimates for F1-score, precision, and recall are presented on the logit scale, while those for FN, FP, and TP are on the log scale. In both scales, a positive estimate indicates a higher mean outcome compared to Spike2, and a negative estimate indicates a lower mean outcome. Significance levels are denoted by asterisks (*, **, ***).

| Model/Outcome metric | F1-Score | Precision | Recall | FN | FP | TP |
| --- | --- | --- | --- | --- | --- | --- |
| One-class SVM | 0.6834 (*) | 0.9705 (*) | -0.20010 | 0.42670 | -1.0024 (*) | -0.12747 |
| SVM | 0.4080 | 0.2958 | 0.09345 | -0.09064 | -1.0309 (*) | -0.08410 |
| XGBoost | 0.9663 (**) | 0.7339 | -0.12733 | 0.17622 | -1.3898 (***) | -0.09989 |

#### Real-world testing: Applying our analysis to a recording with pruritogen injection

Finally, we applied our pipeline under real-world experimental conditions using a pilot example recording obtained during the injection of a chemical itch-inducing substance (BAM 8-22) as previously described [8]. We used the combination of the W_raw_ feature set and XGBoost, selected for its highest F1-score for this specific recording on the background spikes (0.97). The recording characteristics are summarized in Table 8, and F1-scores for all tested combinations are provided in Supplementary Table S6.

**Table 8.**
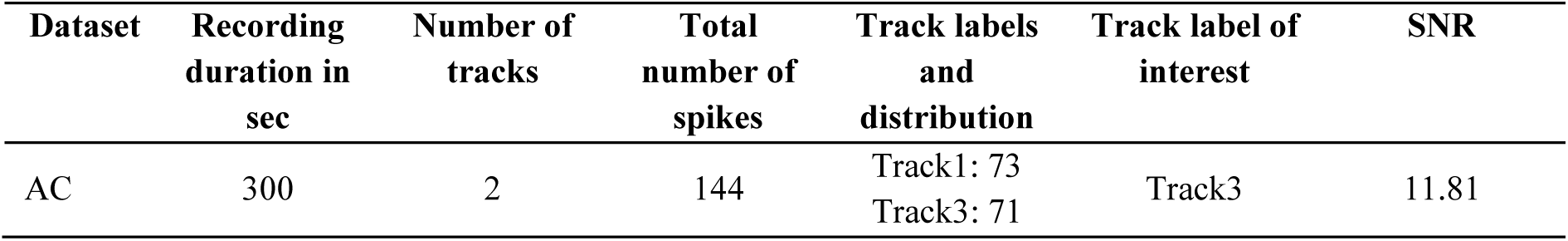
Recording overview with pruritogen injection. The table shows the key characteristics of dataset AC, including the recording duration, the number of tracks, and the total number of extracted spikes. For each track, individual spike counts are listed. As before, we specified the track of interest and the signal-to-noise ratio.

The spike templates for both tracks and a corresponding segment of background noise, along with the template of the fiber of interest and latency, are shown in Figures 11a, 11b, and 11c. Both Spike2 and our pipeline were applied to the same recording. Spike2 assigned a total of 190 spikes to the fiber of interest, while our pipeline assigned 177 spikes. Of these, 174 spikes were assigned by both methods, indicating a high degree of agreement. Three spikes were uniquely assigned by our pipeline, whereas 15 spikes were assigned only by Spike2.

**Figure 11.**
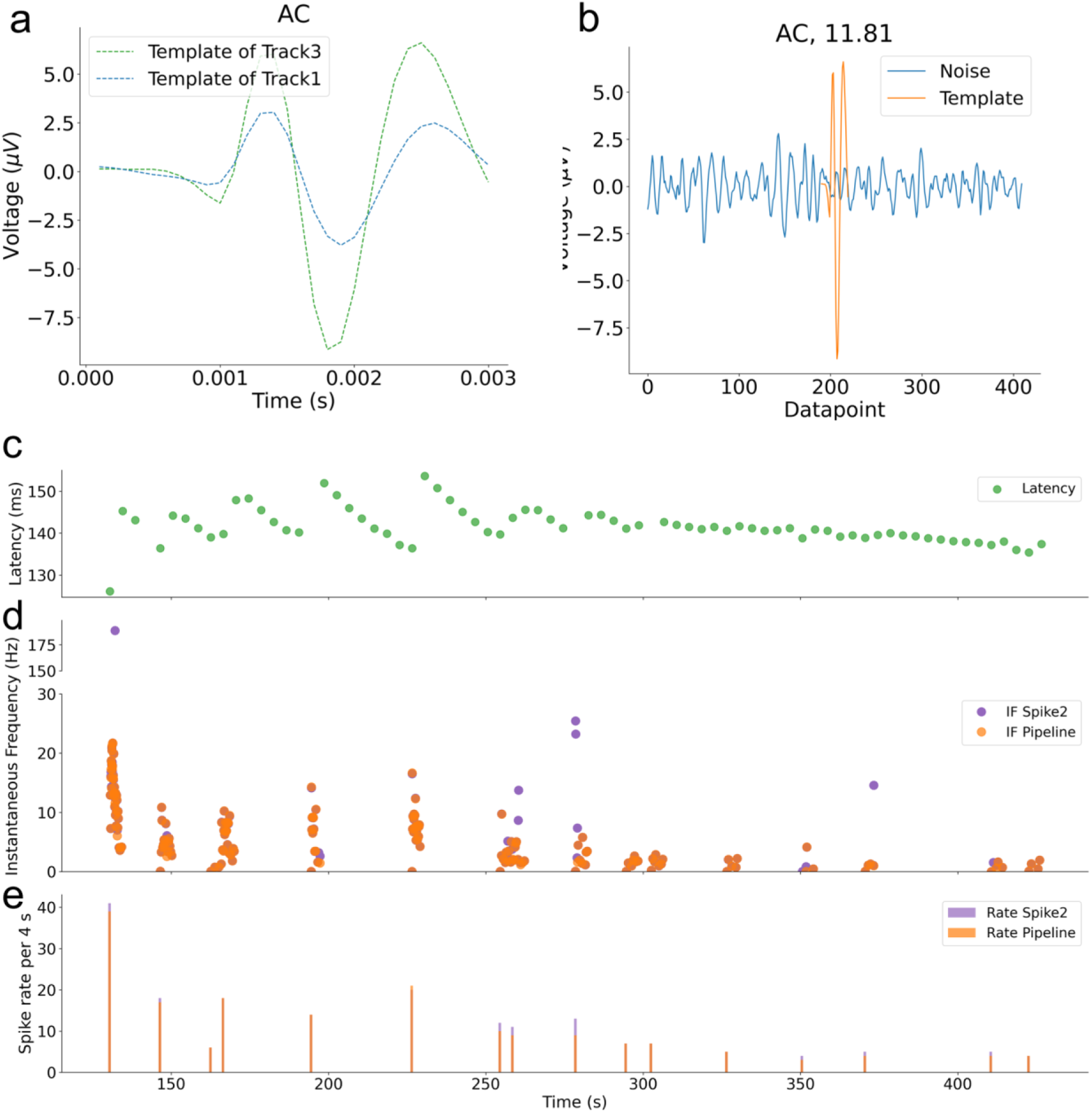
**(a)** Spike template for AC. Average spike template for both tracks, generated by aligning all extracted spikes. This visualization helped to identify the track of interest. **(b)** SNR visualization for AC. We calculated the SNR for the same track template, based on the shown noise segment. For AC, the SNR is 11.81, indicating a very high SNR signal suitable for reliable spike sorting. **(c)** Latency over time for the track of interest. The latency provides critical validation for instantaneous frequency estimates. Sudden shifts in latency reflect the severity of fiber activity. In particular, very high instantaneous frequencies should be accompanied by pronounced latency shifts. **(d)** Instantaneous firing frequency comparison between Spike2 and our pipeline for dataset AC. The instantaneous frequency (IF) was calculated from inter-spike intervals (ISIs) for two consecutive spikes, plotted over time. The purple dots show the IF from spikes sorted with Spike2, which includes at least two high-frequency outliers (200 Hz at 100 seconds, 17 Hz ca. 375 seconds), which could indicate false positives. The orange dots show the frequencies from our pipeline. **(e)** Spike rate comparison between Spike2 and our pipeline for dataset AC. Spike rates were computed by counting spikes within each time interval using the spike sorting results from Spike2 and from our pipeline. The purple dots represent spike rates derived from Spike2, while the orange dots represent spike rates obtained from our pipeline. The two spike rates align across the recording.

To further examine these results, we compared spike shapes and representative signal segments for spikes assigned to the fiber of interest by both methods and for spikes assigned by only one method (see Figure 12). Spikes assigned by both Spike2 and our pipeline exhibited highly similar waveforms and consistent temporal alignment. In contrast, several spikes assigned exclusively by Spike2 occurred in close temporal proximity to neighboring events, indicating implausible discharge frequency and displayed waveform shapes that differed from the dominant template. The mean template of non-agreeing spikes was clearly distinct from those of agreeing spikes, suggesting false assignment rather than activity of the fiber of interest. Representative signal excerpts illustrate that these disagreements typically arise in regions with closely spaced events (see Figure 13).

**Figure 12.**
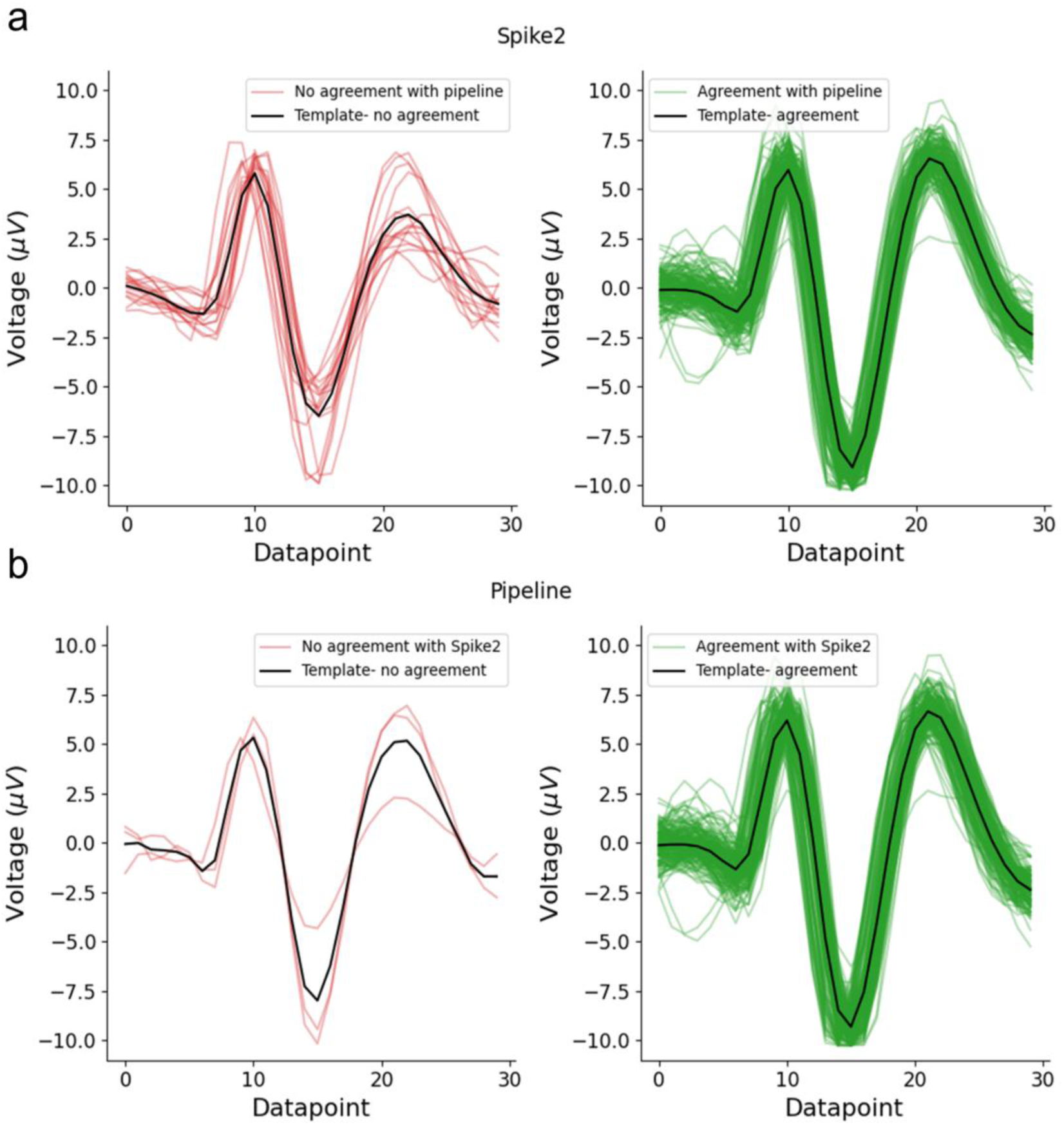
(a) Spike shapes assigned to the fiber of interest by Spike2. Individual spike waveforms are shown, with spikes that agree with the pipeline marked in green and Spike2-only spikes marked in red. The black spike represents the mean waveform (template) computed separately for each category (agreeing and non-agreeing spikes). (b) Same as in (a), but for spikes assigned to the fiber of interest by our pipeline.

**Figure 13.**
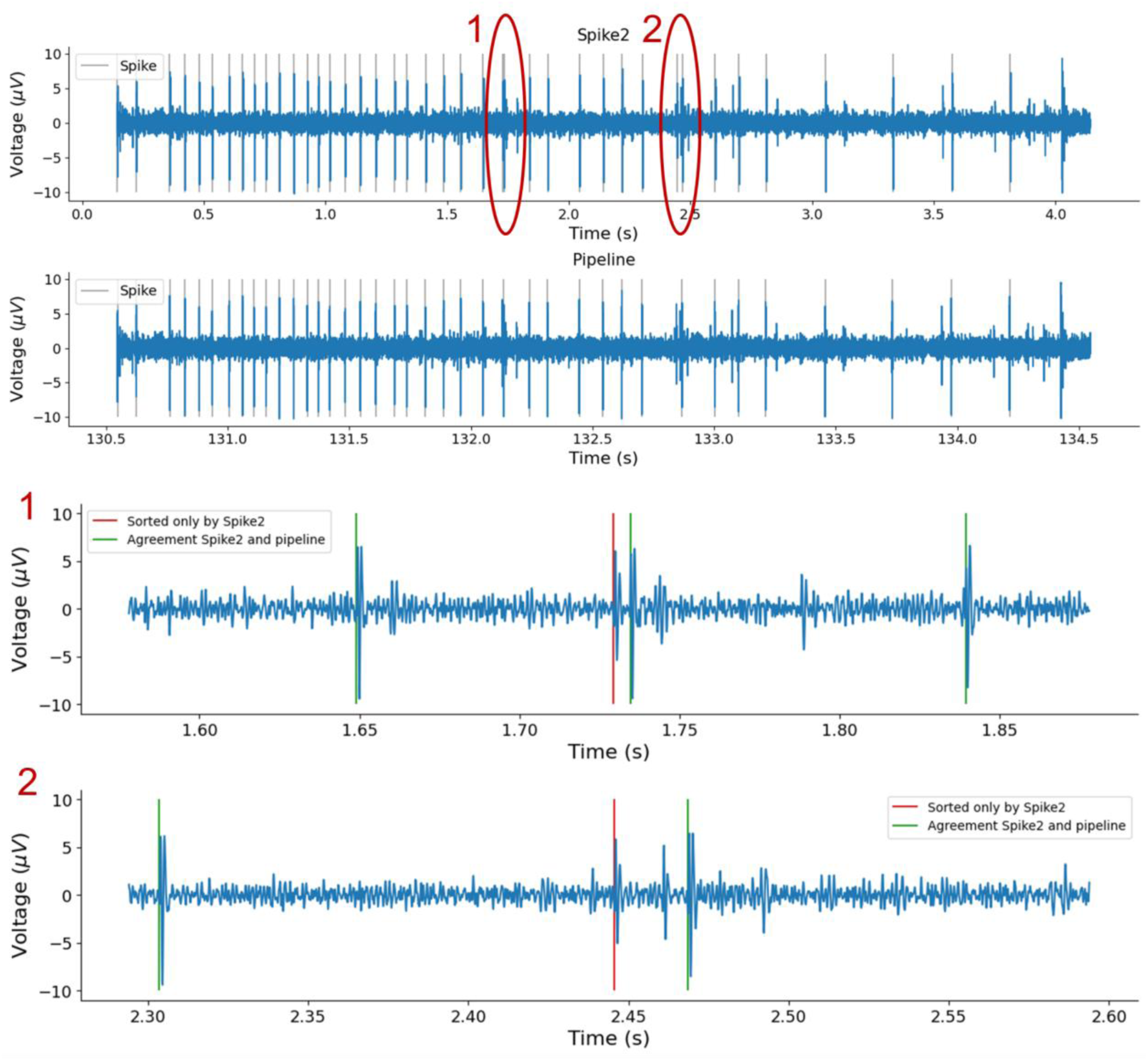
The top panels show the same excerpt of the raw signal analyzed with Spike2 (first panel) and the proposed pipeline (second panel). Detected and sorted spikes assigned to the fiber of interest are indicated by vertical grey lines. Oval markers labeled 1 and 2 denote regions that are shown as zoom-ins in the lower panels. The two bottom panels display the corresponding zoomed signal segments. Spikes assigned by both methods are marked in green, while spikes assigned only by Spike2 are marked in red. The Spike2-only results exhibit lower spike amplitudes and occur in close temporal proximity, illustrating conditions under which a spike belonging to the fiber of interest becomes less certain.

As no ground truth is available, we compared both sorting results indirectly using firing rate metrics. Instantaneous firing frequency (IF) was computed from inter-spike intervals of consecutive spikes (see Figure 11d), and spike rate was additionally analyzed in 4-second bins (see Figure 11e). The mean (± SD) instantaneous firing frequency was 7.9 ± 15.33 Hz for Spike2 and 10.31 ± 40.89 Hz for the pipeline, while the corresponding spike rates were 11.88 ± 9.16 and 11.06 ± 8.98, respectively. In rare cases, Spike2-only assignments resulted in very short inter-spike intervals and correspondingly elevated IF values, including a single instance of approximately 175 Hz arising from two closely spaced spikes. All other Spike2-only spikes exhibited substantially lower firing rates. Together with the shape and signal segment analyses, these results indicate that both methods converge on a largely overlapping set of spikes, while the differences are limited to a small subset of falsely sorted spikes, some of which appear as occasional misassignments in Spike2.

## Discussion

We developed a semi-automated, knowledge- and data-driven supervised spike sorting pipeline for noisy single-electrode microneurography recordings. In the knowledge-driven step, we leverage electrically evoked time-locked spikes obtained via the marking method to limit false detections of spikes from other fibers and to constrain analysis to physiologically relevant signal segments derived due to latency shift and activity-dependent conduction velocity slowing evoked by additional spikes. The same set of time-locked spikes enables the data-driven step by providing labeled training data for supervised spike sorting and recording-specific model selection. The complete pipeline was evaluated on ground truth data created to mimic non-electrically evoked fiber activity. Compared with Spike2, our approach achieved overall higher F1-scores and precision with a relatively smaller loss in recall. On a chemically induced recording, we showed, as proof-of-concept that spikes can be efficiently detected and compared with Spike2, and fewer false-positives are observed.

Simple thresholding yields high recall for spike detection but generates many false positives in recordings with low SNR. More restrictive methods, such as wavelet transforms [14, 32], autoencoders [33], or deep learning-based tools like Variable Projection Networks (VPNets) [34, 35], may enhance precision under challenging conditions. Incorporating physiological constraints, like a maximum expected C-fiber firing rate (<200 Hz) or latency-spike count relationships [36, 37], could further stabilize performance and will be explored in more detail in the future.

High SNR-recordings showed strong classification performance and narrow confidence intervals, whereas recordings with low SNR or multiple active fibers performed poorly, even when template similarity was acceptable. This variability reflects intrinsic challenges of microneurography, where electrode placement, fiber count, and noise vary substantially across experiments.

High variability propagates further to the differences in the best performing feature set-model configurations. Although XGBoost, with the raw waveform feature set, achieved the highest median F1-scores, differences were not statistically significant. Feature importance varied substantially across datasets, indicating that the contribution of individual features depends strongly on the specific recording characteristics. These findings emphasize that spike sorting for microneurography, for now, requires dataset-specific optimization rather than a universal solution.

While our previous work focused on the systematic benchmarking of feature sets for supervised spike sorting using SVMs, the presented work extends this framework by integrating additional classifiers into a complete detection-and-sorting pipeline and by validating performance against ground truth spike timestamps [19]. Although, the earlier work included more recordings from two microneurography laboratories, the lack of ground truth limited evaluation to the tracks only. The availability of verified spike timestamps here, therefore, marks a clear methodological improvement. Consistent with earlier findings, the raw waveform feature set again yields the highest median score. However, the present analysis refines this interpretation by demonstrating that it provides a robust baseline but is not universally optimal, with feature importance varying across recordings.

Several limitations must be considered when interpreting the results. While suitable for electrically evoked responses with clear stimulus timing, chemically induced spike trains may extend across multiple windows and not always exhibit detectable latency shifts within a single interval, particularly at lower firing rates. Future work will therefore focus on more flexible, interval-based strategies within the knowledge-driven framework.

Validation on non-electrically evoked activity is inherently limited by the absence of ground truth. We therefore include only a single recording with chemically induced discharges as a pilot real-world example to demonstrate applicability rather than formal evaluation. Spike2 and our pipeline showed large agreement, with Spike2 identifying 15 additional spikes exhibiting higher instantaneous frequencies and differing spike shapes, which we interpret as misclassified spikes in this example. Without ground truth, however, correctness cannot be established. Agreement between independent methods can only serve as an indicative proxy for reliability in biomedical signal processing [38].

Standard machine learning metrics such as the F1-score can be misleading in physiological applications, as small absolute errors may alter the interpretation of discharge patterns.

Different research questions impose different tolerances: detailed burst analyses, such as patterns within a mechanically evoked burst, require near-perfect accuracy, whereas broader pattern distinctions like the here shown chemical response versus continuous discharges, can tolerate higher error rates. We therefore report additional metrics, such as misassigned spikes per interval, to support application-specific interpretation.

Recordings A4 and A5 showed elevated false-positive rates, which likely reflect genuine spikes from other active fibers, such as sympathetic efferents, which are centrally driven or thermal afferents (e.g., cool-sensitive fibers activated by slight air movement in the room), rather than noise. When waveform shapes are similar, such activity cannot be reliably distinguished without ground truth, representing an inherent limitation of single-electrode recordings.

Overlapping spikes remain a challenge, particularly for spontaneous or chemically evoked activity, and cannot be fully resolved within the current pipeline. However, the experimental design allows partial monitoring of overlap through latency shifts, background activity, and waterfall plots. Latency can support estimating individual fiber activation levels and expected response timing, while tracks and the waterfall plot reveal deviations that can indicate potential overlaps at the time of recording. In rare cases of complete overlap or interference from non-tracked fibers (e.g., non-C-fibers), affected recordings or segments are excluded from further analysis.

Despite the challenges, the overall pipeline performance, combined with the developed open-source workflow, shows good potential in microneurography analysis. The combination of promising results and identified limitations motivates several directions for future development. A primary focus will be the implementation of automated criteria to assess spike sortability based on tracks, template similarity, including signal-to-noise ratio checks, waveform drift monitoring, and comparisons between background stimulation and evoked responses to detect potential response blocks. Automated thresholding and model selection will further reduce the need for manual intervention. The ultimate sortability criteria will additionally require a deeper understanding of how potential errors would affect various spike train characteristics, as commonly used spike train metrics vary substantially in their sensitivity to missing or misassigned spikes [39].

In parallel, we are exploring potentially more generalizable approaches to reduce per-recording retraining by transferring models trained on electrically evoked spikes to other fibers and recordings, while explicitly accounting for inter-fiber variability [40]. Transfer learning strategies may enable more efficient use of large datasets [41].

To address false positives, overlapping spikes and the sorting verification of chemically or spontaneously evoked activity, we are developing a post-sorting verification stage based on latency prediction. This approach estimates expected spike counts and latencies and compares them to observed values, allowing the identification of physiologically improbable label assignments [36, 37].

Finally, we plan a systematic evaluation across recordings with chemically, mechanically, and thermally evoked spikes, exploring the potential of our pipeline to enable reliable, reusable, and computationally efficient analysis of evoked activity of C-fibers, and, in a long-term perspective, the analysis of spontaneous nociceptor activity in neuropathic pain.

## Acknowledgements

We acknowledge Danxia Bao for her technical support.

## Author contributions

B.N. and E.K. conceptualized the work; B.N., A.F., and A.M. performed the experiments and acquired the data; A.F. curated the data; A.T., E.K., B.N., and A.M. analyzed the data; A.T. developed the computational pipeline; E.K. and B.N. interpreted the data; All authors critically discussed the results; A.T. wrote the main manuscript text and prepared the figures; All authors have revised the manuscript and approved the final version.

## Data availability statement

The raw datasets generated and/or analyzed during the current work are available from the corresponding author on reasonable request. The source code and selected test data are available on https://github.com/Digital-C-Fiber/SpikeSortingForSpikingPatterns.

