## Supplementary Material for "A Hybrid Knowledge- and Data-driven Model for Automatic Assessment of Induced Spiking Patterns in C-fiber Microneurography: Methodological Development"

### Supplementary Methods

To quantify sorting performance, we computed standard classification metrics based on the number of true positives (TP), false positives (FP), and false negatives (FN). Precision and recall were defined as:

$$\text{Precision} = \frac{\text{TP}}{\text{TP} + \text{FP}}, \text{Recall} = \frac{\text{TP}}{\text{TP} + \text{FN}}.$$

From these, the F1-score was obtained as:

$$\text{F1-score} = \frac{2 \times \text{Precision} \times \text{Recall}}{\text{Precision} \times \text{Recall}}$$

The False Discovery Rate (FDR) was computed as:

$$\text{FDR} = \frac{\text{FP}}{\text{TP} + \text{FP}} = 1 - \text{Precision}.$$

Because the detection and sorting were performed on 4-s background stimulus-locked intervals, we additionally computed the number of incorrect assignments per interval:

$$\text{FP per interval} = \frac{\text{FP}}{N_{\text{intervals}}},$$

where $N_{\text{intervals}}$ is the number of stimulation intervals searched for each recording.

To characterize template similarity and distance, we also report the mean squared error (MSE), root mean squared error (RMSE), and mean absolute error (MAE) between the template of the fiber of interest $x$ and other templates $y$:

$$\text{MSE} = \frac{1}{N}\sum_{i=1}^{N}(x_i - y_i)^2, \text{RMSE} = \sqrt{\text{MSE}}, \text{MAE} = \frac{1}{N}\sum_{i=1}^{N} | x_i - y_i |.$$

### Supplementary Figures

**Figure S1. Stimulation protocols for ground truth recordings.** For each ground truth recording, the applied stimulation protocol is shown together with the latencies of tracked spikes. Green dots indicate the latencies of individual tracked spikes. The red dashed line denotes the constant background electrical stimulation frequency (0.25 Hz). Colored circles indicate the number of pulses of additional stimulation, a single additional pulse is marked by a cross. The height of the lines attached to the colored circles indicates the frequency of the additional stimulation, while their color encodes the temporal distance between the last additional pulse and the background stimulus. A detailed description of the stimulation protocol visualization can be found in Kutafina et al., 2022.

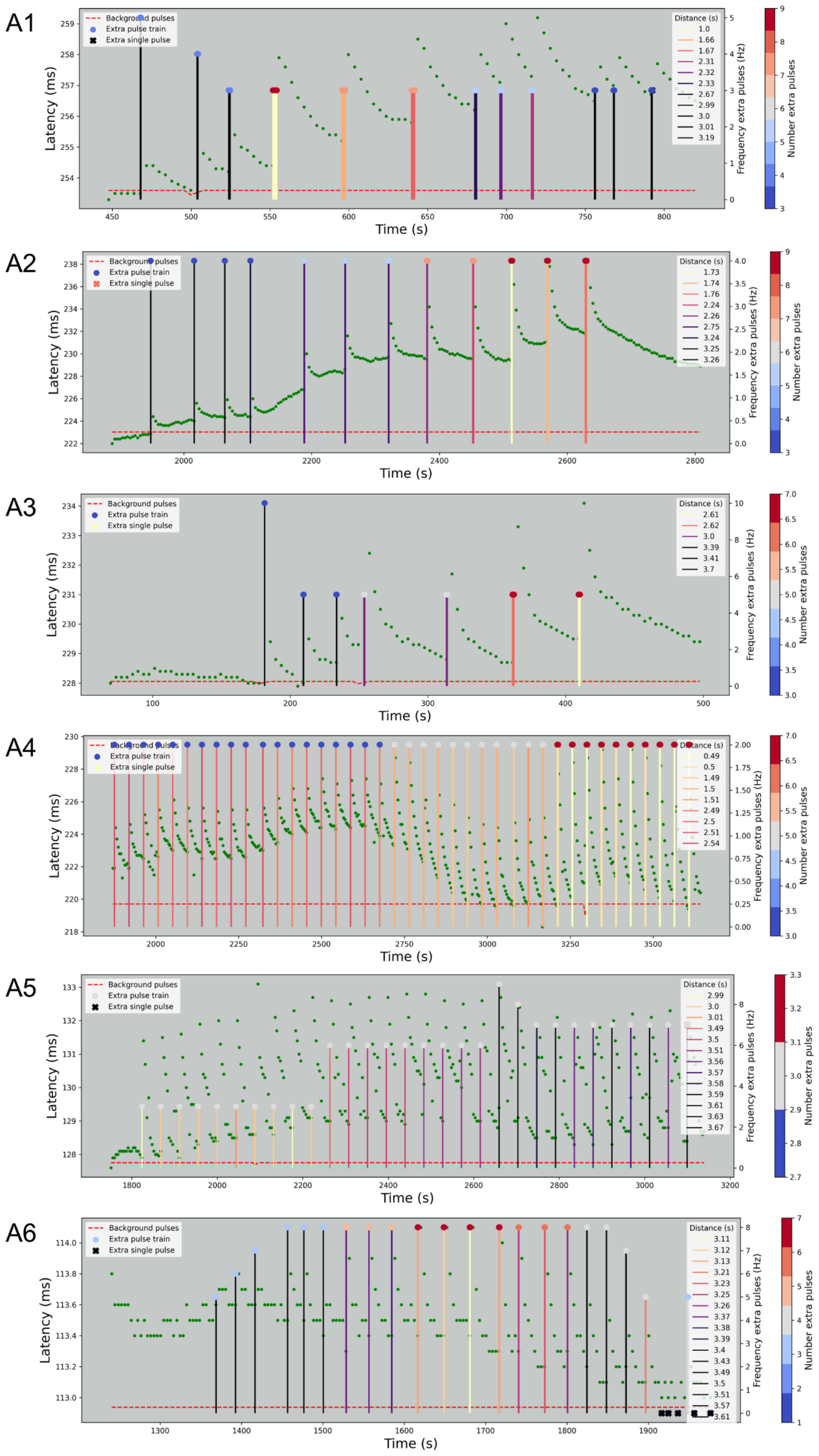

A1
Background pulses
Extra pulse train
Extra single pulse
Distance (s)
1.0
1.66
1.67
2.31
2.32
2.33
2.67
2.99
3.0
3.01
3.19
Latency (ms)
Time (s)
Frequency extra pulses (Hz)
Number extra pulses
A2
Background pulses
Extra pulse train
Extra single pulse
Distance (s)
1.73
1.74
1.76
2.24
2.26
2.75
3.24
3.25
3.26
Latency (ms)
Time (s)
Frequency extra pulses (Hz)
Number extra pulses
A3
Background pulses
Extra pulse train
Extra single pulse
Distance (s)
2.61
2.62
3.0
3.39
3.41
3.7
Latency (ms)
Time (s)
Frequency extra pulses (Hz)
Number extra pulses
A4
Background pulses
Extra pulse train
Extra single pulse
Distance (s)
0.49
0.5
1.49
1.5
1.51
2.49
2.5
2.51
2.54
Latency (ms)
Time (s)
Frequency extra pulses (Hz)
Number extra pulses
A5
Background pulses
Extra pulse train
Extra single pulse
Distance (s)
2.99
3.0
3.01
3.49
3.5
3.51
3.56
3.57
3.58
3.59
3.61
3.63
3.67
Latency (ms)
Time (s)
Frequency extra pulses (Hz)
Number extra pulses
A6
Background pulses
Extra pulse train
Extra single pulse
Distance (s)
3.11
3.12
3.13
3.21
3.23
3.25
3.26
3.37
3.38
3.39
3.4
3.43
3.49
3.5
3.51
3.57
3.61
Latency (ms)
Time (s)
Frequency extra pulses (Hz)
Number extra pulses

**Figure S2**. **Waveform drift of the fiber of interest templates over time across all recordings.** For each recording, spike templates were computed separately for the beginning, middle, and end of the recording by averaging all tracked spikes within each respective segment. Templates derived from the beginning are shown in green, those from the middle in dashed red, and those from the end in dashed blue.

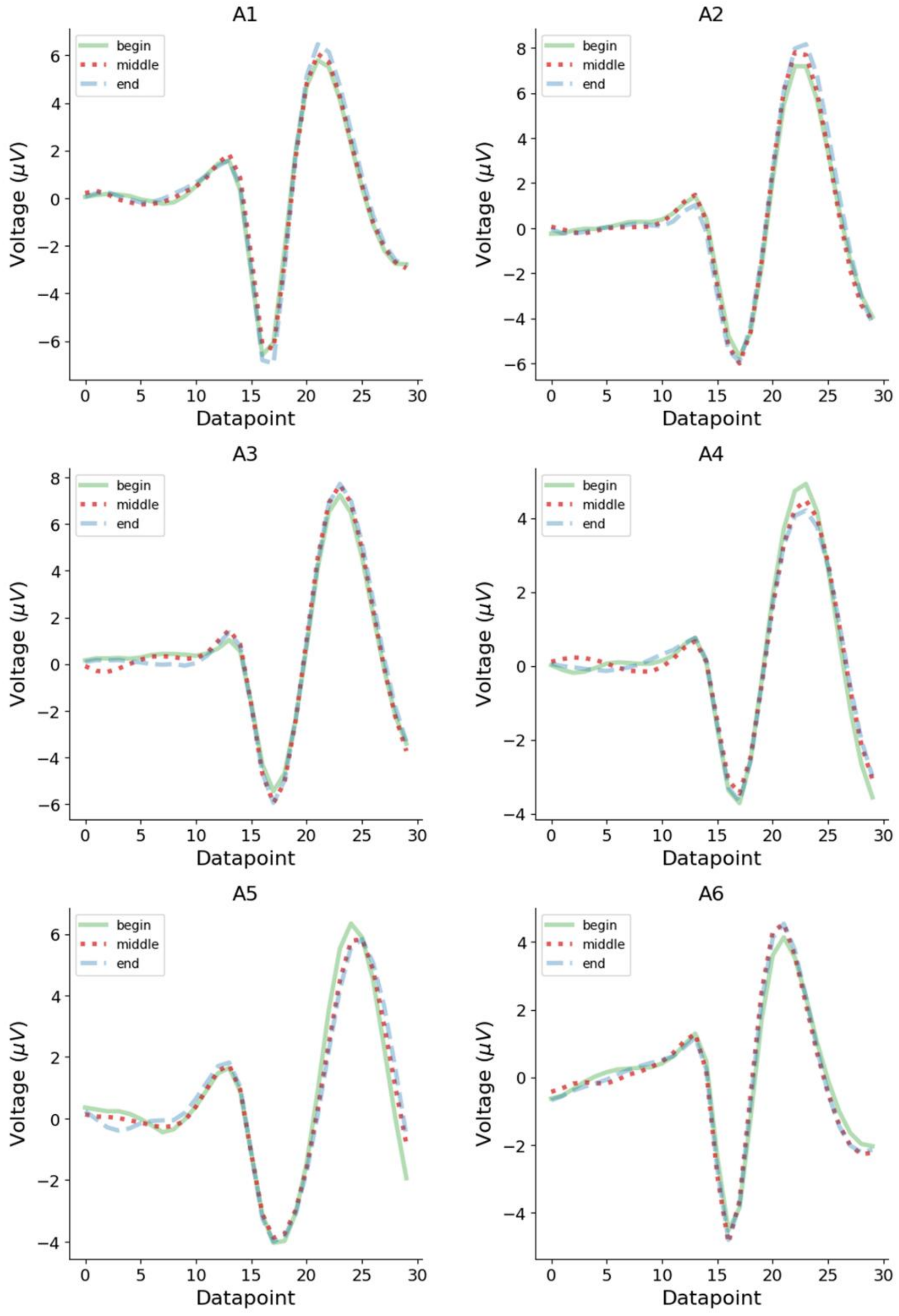


---

**Figure S3. Feature importance analysis across feature sets and models**. Feature importance rankings for the ten most informative features are shown for three feature sets: **(a)** SPDF, **(b)** $SPDF_{FV3}$, and **(c)** $W_{raw}$. For each panel, results obtained with a random forest classifier are shown in the left column, while results from an L1-regularized logistic regression model are shown in the right column. Importance scores were computed separately for each recording and model to identify features that contributed most strongly to spike sorting performance.

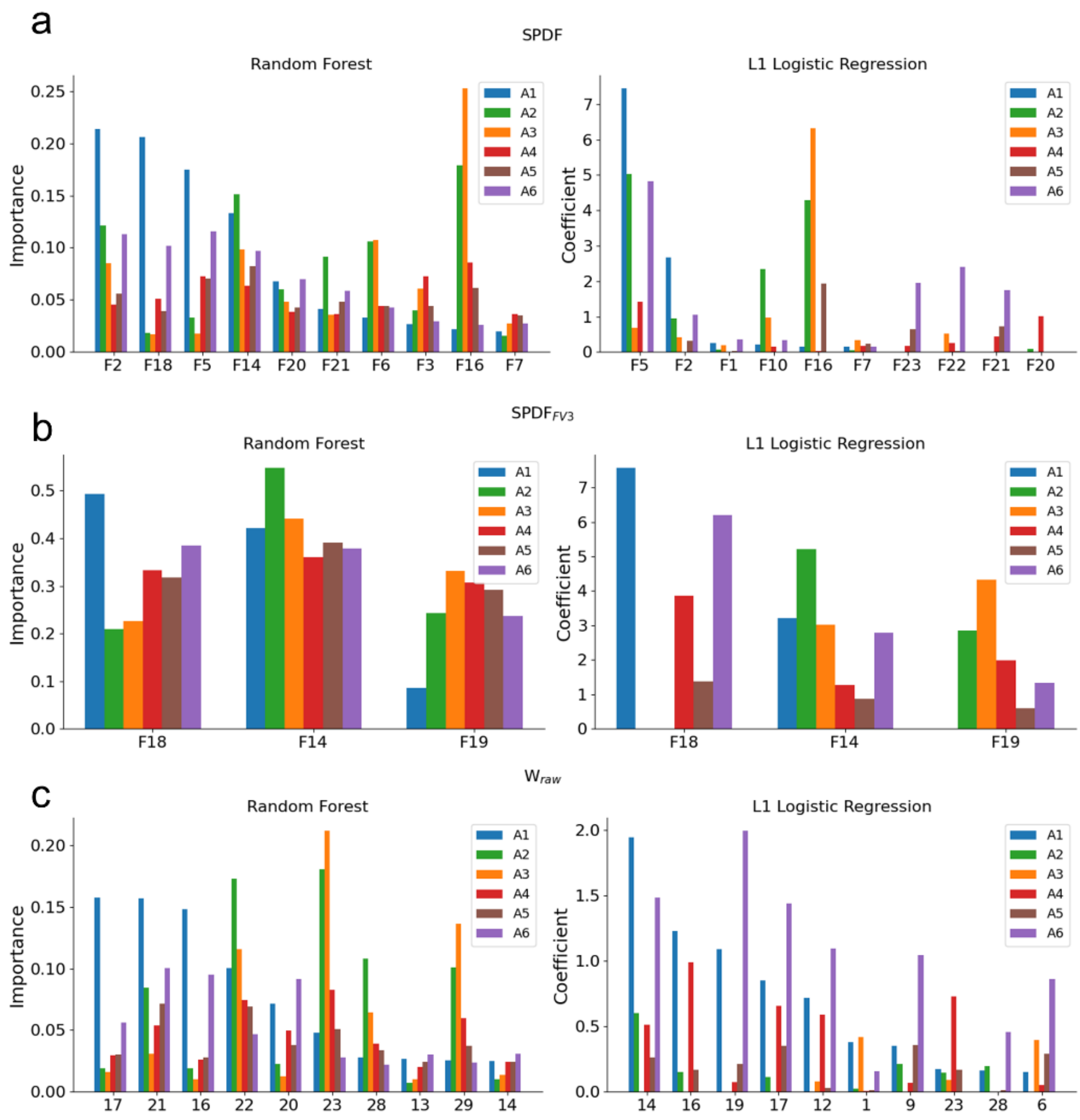


### Supplementary Tables

**Table S1**. **Performance metrics for spike detection using thresholding.** Detection results are summarized by the number of true positives (TP), false negatives (FN), and false positives (FP), together with the derived evaluation metrics precision, recall, and F1-score.

| Dataset | Detected spike count | TP | FN | FP | F1-Score | Precision | Recall |
|---|---|---|---|---|---|---|---|
| A1 | 598 | 64 | 1 | 534 | 0.19 | 0.11 | 0.98 |
| A2 | 357 | 80 | 0 | 277 | 0.37 | 0.22 | 1.0 |
| A3 | 92 | 30 | 1 | 62 | 0.49 | 0.33 | 0.97 |
| A4 | 4643 | 192 | 22 | 4451 | 0.08 | 0.04 | 0.9 |

| | | | | | | | |
|---|---|---|---|---|---|---|---|
| A5 | 1802 | 109 | 6 | 1693 | 0.11 | 0.06 | 0.95 |
| A6 | 497 | 48 | 4 | 449 | 0.17 | 0.1 | 0.92 |

**Table S2. Summary of spike sorting performance across models and feature sets for our pipeline.** Spike sorting results are reported for the different classification models and feature sets, including true positives (TP), false positives (FP), false negatives (FN), and true negatives (TN), along with the corresponding precision, recall, and F1-score.

| Dataset | Model | Feature Set | TP | FP | TN | FN | F1-Score | Precision | Recall |
|---|---|---|---|---|---|---|---|---|---|
| A1 | svm | spdf | 64.0 | 4.0 | 520.0 | 0.0 | 0.97 | 0.94 | 1.0 |
| A2 | svm | spdf | 76.0 | 32.0 | 236.0 | 4.0 | 0.81 | 0.7 | 0.95 |
| A3 | svm | spdf | 28.0 | 24.0 | 38.0 | 2.0 | 0.68 | 0.54 | 0.93 |
| A4 | svm | spdf | 190.0 | 4355.0 | 5.0 | 0.0 | 0.08 | 0.04 | 1.0 |
| A5 | svm | spdf | 85.0 | 576.0 | 1063.0 | 23.0 | 0.22 | 0.13 | 0.79 |
| A6 | svm | spdf | 44.0 | 86.0 | 356.0 | 4.0 | 0.49 | 0.34 | 0.92 |
| A1 | svm | spdf_fv3 | 63.0 | 5.0 | 519.0 | 1.0 | 0.95 | 0.93 | 0.98 |
| A2 | svm | spdf_fv3 | 76.0 | 44.0 | 224.0 | 4.0 | 0.76 | 0.63 | 0.95 |
| A3 | svm | spdf_fv3 | 27.0 | 24.0 | 38.0 | 3.0 | 0.67 | 0.53 | 0.9 |
| A4 | svm | spdf_fv3 | 190.0 | 4360.0 | 0.0 | 0.0 | 0.08 | 0.04 | 1.0 |
| A5 | svm | spdf_fv3 | 0.0 | 0.0 | 1639.0 | 108.0 | 0.0 | 0.0 | 0.0 |
| A6 | svm | spdf_fv3 | 44.0 | 46.0 | 396.0 | 4.0 | 0.64 | 0.49 | 0.92 |
| A1 | svm | w_raw | 56.0 | 3.0 | 531.0 | 8.0 | 0.91 | 0.95 | 0.88 |
| A2 | svm | w_raw | 77.0 | 47.0 | 230.0 | 3.0 | 0.75 | 0.62 | 0.96 |
| A3 | svm | w_raw | 29.0 | 12.0 | 50.0 | 1.0 | 0.82 | 0.71 | 0.97 |
| A4 | svm | w_raw | 148.0 | 1237.0 | 3214.0 | 44.0 | 0.19 | 0.11 | 0.77 |
| A5 | svm | w_raw | 88.0 | 126.0 | 1567.0 | 21.0 | 0.54 | 0.41 | 0.81 |
| A6 | svm | w_raw | 43.0 | 73.0 | 376.0 | 5.0 | 0.52 | 0.37 | 0.9 |
| A1 | one_class_svm | spdf | 57.0 | 4.0 | 520.0 | 7.0 | 0.91 | 0.93 | 0.89 |
| A2 | one_class_svm | spdf | 58.0 | 52.0 | 216.0 | 22.0 | 0.61 | 0.53 | 0.72 |
| A3 | one_class_svm | spdf | 25.0 | 27.0 | 35.0 | 5.0 | 0.61 | 0.48 | 0.83 |
| A4 | one_class_svm | spdf | 84.0 | 180.0 | 4180.0 | 106.0 | 0.37 | 0.32 | 0.44 |
| A5 | one_class_svm | spdf | 88.0 | 791.0 | 848.0 | 20.0 | 0.18 | 0.1 | 0.81 |

| | | | | | | | | | |
|---|---|---|---|---|---|---|---|---|---|
| A6 | one_class_svm | spdf | 29.0 | 9.0 | 433.0 | 19.0 | 0.67 | 0.76 | 0.6 |
| A1 | one_class_svm | spdf_fv3 | 63.0 | 4.0 | 520.0 | 1.0 | 0.96 | 0.94 | 0.98 |
| A2 | one_class_svm | spdf_fv3 | 77.0 | 55.0 | 213.0 | 3.0 | 0.73 | 0.58 | 0.96 |
| A3 | one_class_svm | spdf_fv3 | 25.0 | 24.0 | 38.0 | 5.0 | 0.63 | 0.51 | 0.83 |
| A4 | one_class_svm | spdf_fv3 | 178.0 | 3223.0 | 1137.0 | 12.0 | 0.1 | 0.05 | 0.94 |
| A5 | one_class_svm | spdf_fv3 | 95.0 | 1134.0 | 505.0 | 13.0 | 0.14 | 0.08 | 0.88 |
| A6 | one_class_svm | spdf_fv3 | 44.0 | 54.0 | 388.0 | 4.0 | 0.6 | 0.45 | 0.92 |
| A1 | one_class_svm | w_raw | 55.0 | 7.0 | 527.0 | 9.0 | 0.87 | 0.89 | 0.86 |
| A2 | one_class_svm | w_raw | 74.0 | 19.0 | 258.0 | 6.0 | 0.86 | 0.8 | 0.92 |
| A3 | one_class_svm | w_raw | 27.0 | 11.0 | 51.0 | 3.0 | 0.79 | 0.71 | 0.9 |
| A4 | one_class_svm | w_raw | 175.0 | 2198.0 | 2253.0 | 17.0 | 0.14 | 0.07 | 0.91 |
| A5 | one_class_svm | w_raw | 47.0 | 0.0 | 1693.0 | 62.0 | 0.6 | 1.0 | 0.43 |
| A6 | one_class_svm | w_raw | 46.0 | 104.0 | 345.0 | 2.0 | 0.46 | 0.31 | 0.96 |
| A1 | xgboost | spdf | 62.0 | 2.0 | 522.0 | 2.0 | 0.97 | 0.97 | 0.97 |
| A2 | xgboost | spdf | 76.0 | 29.0 | 239.0 | 4.0 | 0.82 | 0.72 | 0.95 |
| A3 | xgboost | spdf | 27.0 | 34.0 | 28.0 | 3.0 | 0.59 | 0.44 | 0.9 |
| A4 | xgboost | spdf | 141.0 | 1292.0 | 3068.0 | 49.0 | 0.17 | 0.1 | 0.74 |
| A5 | xgboost | spdf | 78.0 | 236.0 | 1403.0 | 30.0 | 0.37 | 0.25 | 0.72 |
| A6 | xgboost | spdf | 44.0 | 21.0 | 421.0 | 4.0 | 0.78 | 0.68 | 0.92 |
| A1 | xgboost | spdf_fv3 | 62.0 | 3.0 | 521.0 | 2.0 | 0.96 | 0.95 | 0.97 |
| A2 | xgboost | spdf_fv3 | 76.0 | 49.0 | 219.0 | 4.0 | 0.74 | 0.61 | 0.95 |
| A3 | xgboost | spdf_fv3 | 26.0 | 21.0 | 41.0 | 4.0 | 0.68 | 0.55 | 0.87 |
| A4 | xgboost | spdf_fv3 | 122.0 | 1291.0 | 3069.0 | 68.0 | 0.15 | 0.09 | 0.64 |
| A5 | xgboost | spdf_fv3 | 66.0 | 335.0 | 1304.0 | 42.0 | 0.26 | 0.16 | 0.61 |
| A6 | xgboost | spdf_fv3 | 44.0 | 41.0 | 401.0 | 4.0 | 0.66 | 0.52 | 0.92 |
| A1 | xgboost | w_raw | 52.0 | 9.0 | 525.0 | 12.0 | 0.83 | 0.85 | 0.81 |
| A2 | xgboost | w_raw | 76.0 | 21.0 | 256.0 | 4.0 | 0.86 | 0.78 | 0.95 |
| A3 | xgboost | w_raw | 29.0 | 10.0 | 52.0 | 1.0 | 0.84 | 0.74 | 0.97 |
| A4 | xgboost | w_raw | 148.0 | 1284.0 | 3167.0 | 44.0 | 0.18 | 0.1 | 0.77 |
| A5 | xgboost | w_raw | 90.0 | 146.0 | 1547.0 | 19.0 | 0.52 | 0.38 | 0.83 |

| | | | | | | | | | |
|---|---|---|---|---|---|---|---|---|---|
| A6 | xgboost | w_raw | 40.0 | 27.0 | 422.0 | 8.0 | 0.7 | 0.6 | 0.83 |

**Table S3. Best spike sorting performance per dataset.** For each recording, the classifier-feature set combination yielding the highest F1-score was selected. Results are summarized as true positives (TP), false positives (FP), true negatives (TN), and false negatives (FN), together with precision, recall, F1-score, false discovery rate (FDR), the number of analyzed 4-second intervals, and the mean number of false positives per interval.

| Dataset | TP | FP | TN | FN | F1-Score | Precision | Recall | FDR | Number of Intervals | FP per Interval |
|---|---|---|---|---|---|---|---|---|---|---|
| A1 | 64.0 | 4.0 | 520.0 | 0.0 | 0.97 | 0.94 | 1.0 | 0.06 | 11 | 0.36 |
| A2 | 76.0 | 21.0 | 256.0 | 4.0 | 0.86 | 0.78 | 0.95 | 0.20 | 12 | 1.75 |
| A3 | 29.0 | 10.0 | 52.0 | 1.0 | 0.84 | 0.74 | 0.97 | 0.26 | 6 | 1.67 |
| A4 | 84.0 | 180.0 | 4180.0 | 106.0 | 0.37 | 0.32 | 0.44 | 0.68 | 39 | 4.62 |
| A5 | 47.0 | 0.0 | 1693.0 | 62.0 | 0.6 | 1.0 | 0.43 | 0 | 29 | 0 |
| A6 | 44.0 | 21.0 | 421.0 | 4.0 | 0.78 | 0.68 | 0.92 | 0.32 | 21 | 1.0 |

**Table S4. Detection and sorting performance using Spike2.** Spike detection and sorting results obtained with Spike2 on the full ground truth datasets are summarized by the number of true positives (TP), false positives (FP), and false negatives (FN), together with the derived precision, recall, and F1-score.

| Dataset | FP | TP | FN | F1-Score | Precision | Recall |
|---|---|---|---|---|---|---|
| A1 | 20.0 | 140.0 | 8.0 | 0.91 | 0.88 | 0.95 |
| A2 | 782.0 | 271.0 | 28.0 | 0.4 | 0.26 | 0.91 |
| A3 | 654.0 | 164.0 | 9.0 | 0.33 | 0.2 | 0.95 |
| A4 | 20761.0 | 472.0 | 153.0 | 0.04 | 0.02 | 0.76 |
| A5 | 24494.0 | 398.0 | 38.0 | 0.03 | 0.02 | 0.91 |
| A6 | 1447.0 | 262.0 | 26.0 | 0.26 | 0.15 | 0.91 |

**Table S5. Detection and sorting performance using Spike2 within the constrained post-hoc search space.** Spike detection and sorting results obtained with Spike2 are evaluated only on intervals included in the constrained post-hoc search space used for comparison with our pipeline. Performance is reported in terms of true positives (TP), false positives (FP), and false negatives (FN), together with the corresponding precision, recall, and F1-score.

| Dataset | FP | TP | FN | F1-Score | Precision | Recall |
|---|---|---|---|---|---|---|
| A1 | 5.0 | 61.0 | 4.0 | 0.93 | 0.92 | 0.94 |
| A2 | 78.0 | 76.0 | 4.0 | 0.65 | 0.49 | 0.95 |
| A3 | 71.0 | 29.0 | 2.0 | 0.44 | 0.29 | 0.94 |
| A4 | 1944.0 | 159.0 | 55.0 | 0.14 | 0.08 | 0.74 |

| | | | | | | |
|---|---|---|---|---|---|---|
| A5 | 2991.0 | 96.0 | 19.0 | 0.06 | 0.03 | 0.83 |
| A6 | 186.0 | 48.0 | 4.0 | 0.34 | 0.21 | 0.92 |

**Table S6. Model and feature set comparison for dataset AC.** Average F1-scores from 5-fold cross-validation on background spikes are reported for all model-feature set combinations with evaluation restricted to tracked spikes to identify the best-performing model-feature set combination in the absence of ground truth.

| Dataset | Model | Feature Set | F1-Score |
|---|---|---|---|
| AC | svm | spdf | 0.92 |
| AC | svm | spdf_fv3 | 0.75 |
| AC | svm | w_raw | 0.85 |
| AC | one_class_svm | spdf | 0.96 |
| AC | one_class_svm | spdf_fv3 | 0.95 |
| AC | one_class_svm | w_raw | 0.97 |
| AC | xgboost | spdf | 0.97 |
| AC | xgboost | spdf_fv3 | 0.95 |
| AC | xgboost | w_raw | 0.97 |

**Table S7. Spike characteristics for dataset AC detected and sorted using Spike2.** Detailed spike information is reported for dataset AC following detection and sorting with Spike2, including spike identifiers, timestamps, inter-spike intervals (ISI), and instantaneous firing frequencies (Hz).

| Spike ID | Spike Timestamp | Inter-spike-interval (ISI) | Instantaneous Frequency (Hz) |
|---|---|---|---|
| 0 | 130.54 | | |
| 1 | 130.62 | 0.08 | 12.92 |
| 2 | 130.76 | 0.14 | 7.27 |
| 3 | 130.82 | 0.06 | 16.13 |
| 4 | 130.88 | 0.06 | 16.84 |
| 5 | 130.93 | 0.05 | 18.69 |
| 6 | 131.0 | 0.07 | 14.33 |
| 7 | 131.06 | 0.06 | 18.08 |
| 8 | 131.11 | 0.05 | 20.88 |
| 9 | 131.15 | 0.05 | 20.45 |
| 10 | 131.21 | 0.05 | 18.48 |
| 11 | 131.27 | 0.06 | 16.92 |
| 12 | 131.32 | 0.06 | 17.92 |
| 13 | 131.37 | 0.05 | 21.55 |
| 14 | 131.42 | 0.05 | 21.69 |
| 15 | 131.48 | 0.06 | 15.7 |
| 16 | 131.54 | 0.06 | 16.31 |
| 17 | 131.6 | 0.06 | 15.5 |
| 18 | 131.68 | 0.08 | 12.82 |
| 19 | 131.73 | 0.05 | 19.92 |
| 20 | 131.81 | 0.08 | 12.94 |
| 21 | 131.88 | 0.07 | 13.5 |

| | | | |
|---|---|---|---|
| 22 | 131.95 | 0.07 | 14.29 |
| 23 | 132.05 | 0.09 | 10.95 |
| 24 | 132.13 | 0.08 | 12.42 |
| 25 | 132.13 | 0.01 | 188.68 |
| 26 | 132.24 | 0.1 | 9.55 |
| 27 | 132.31 | 0.08 | 13.25 |
| 28 | 132.44 | 0.13 | 7.7 |
| 29 | 132.54 | 0.1 | 10.16 |
| 30 | 132.62 | 0.08 | 12.92 |
| 31 | 132.7 | 0.08 | 12.06 |
| 32 | 132.84 | 0.14 | 7.04 |
| 33 | 132.87 | 0.02 | 43.48 |
| 34 | 133.0 | 0.13 | 7.42 |
| 35 | 133.1 | 0.1 | 10.21 |
| 36 | 133.21 | 0.11 | 8.98 |
| 37 | 133.45 | 0.24 | 4.09 |
| 38 | 133.73 | 0.28 | 3.62 |
| 39 | 133.97 | 0.24 | 4.12 |
| 40 | 134.21 | 0.24 | 4.17 |
| 41 | 146.55 | 12.34 | 0.08 |
| 42 | 146.85 | 0.3 | 3.32 |
| 43 | 146.95 | 0.09 | 10.85 |
| 44 | 147.06 | 0.12 | 8.69 |
| 45 | 147.31 | 0.24 | 4.08 |
| 46 | 147.52 | 0.22 | 4.56 |
| 47 | 147.71 | 0.19 | 5.38 |
| 48 | 147.95 | 0.23 | 4.26 |
| 49 | 148.14 | 0.19 | 5.27 |
| 50 | 148.26 | 0.12 | 8.12 |
| 51 | 148.48 | 0.22 | 4.44 |
| 52 | 148.65 | 0.16 | 6.09 |
| 53 | 148.94 | 0.3 | 3.37 |
| 54 | 149.14 | 0.19 | 5.13 |
| 55 | 149.32 | 0.18 | 5.65 |
| 56 | 149.55 | 0.23 | 4.33 |
| 57 | 149.84 | 0.3 | 3.38 |
| 58 | 150.21 | 0.37 | 2.71 |
| 59 | 162.55 | 12.34 | 0.08 |
| 60 | 163.83 | 1.28 | 0.78 |
| 61 | 165.12 | 1.28 | 0.78 |
| 62 | 165.88 | 0.76 | 1.31 |
| 63 | 166.0 | 0.12 | 8.36 |
| 64 | 166.27 | 0.27 | 3.69 |
| 65 | 166.56 | 0.28 | 3.54 |
| 66 | 166.7 | 0.14 | 6.97 |
| 67 | 166.8 | 0.1 | 10.26 |
| 68 | 166.96 | 0.16 | 6.28 |
| 69 | 167.1 | 0.15 | 6.85 |
| 70 | 167.22 | 0.12 | 8.29 |
| 71 | 167.37 | 0.15 | 6.8 |
| 72 | 167.51 | 0.14 | 6.9 |
| 73 | 167.64 | 0.12 | 8.19 |
| 74 | 167.86 | 0.22 | 4.57 |
| 75 | 167.98 | 0.12 | 8.29 |
| 76 | 168.1 | 0.12 | 8.11 |
| 77 | 168.42 | 0.32 | 3.1 |
| 78 | 168.53 | 0.11 | 9.38 |
| 79 | 168.88 | 0.35 | 2.85 |

| | | | |
|---|---|---|---|
| 80 | 169.43 | 0.55 | 1.82 |
| 81 | 169.69 | 0.26 | 3.9 |
| 82 | 169.98 | 0.29 | 3.47 |
| 83 | 194.45 | 24.47 | 0.04 |
| 84 | 194.52 | 0.07 | 14.16 |
| 85 | 194.66 | 0.14 | 7.1 |
| 86 | 194.77 | 0.11 | 9.12 |
| 87 | 194.92 | 0.15 | 6.55 |
| 88 | 195.03 | 0.11 | 9.12 |
| 89 | 195.14 | 0.11 | 9.12 |
| 90 | 195.28 | 0.14 | 7.23 |
| 91 | 195.57 | 0.29 | 3.45 |
| 92 | 195.86 | 0.29 | 3.47 |
| 93 | 195.95 | 0.1 | 10.48 |
| 94 | 196.57 | 0.61 | 1.63 |
| 95 | 196.88 | 0.31 | 3.22 |
| 96 | 197.26 | 0.38 | 2.62 |
| 97 | 226.49 | 29.23 | 0.03 |
| 98 | 226.55 | 0.06 | 16.53 |
| 99 | 226.69 | 0.14 | 7.2 |
| 100 | 226.8 | 0.11 | 9.43 |
| 101 | 226.95 | 0.15 | 6.72 |
| 102 | 227.06 | 0.11 | 8.84 |
| 103 | 227.16 | 0.1 | 9.73 |
| 104 | 227.3 | 0.14 | 7.29 |
| 105 | 227.4 | 0.1 | 9.55 |
| 106 | 227.54 | 0.13 | 7.55 |
| 107 | 227.62 | 0.08 | 12.39 |
| 108 | 227.8 | 0.19 | 5.32 |
| 109 | 227.95 | 0.15 | 6.78 |
| 110 | 228.08 | 0.13 | 7.54 |
| 111 | 228.27 | 0.18 | 5.42 |
| 112 | 228.4 | 0.13 | 7.76 |
| 113 | 228.57 | 0.17 | 5.96 |
| 114 | 228.7 | 0.13 | 7.62 |
| 115 | 228.82 | 0.13 | 7.89 |
| 116 | 229.06 | 0.23 | 4.28 |
| 117 | 254.56 | 25.5 | 0.04 |
| 118 | 254.92 | 0.37 | 2.71 |
| 119 | 255.03 | 0.1 | 9.75 |
| 120 | 255.49 | 0.47 | 2.15 |
| 121 | 256.14 | 0.65 | 1.55 |
| 122 | 256.44 | 0.3 | 3.29 |
| 123 | 256.8 | 0.36 | 2.79 |
| 124 | 256.99 | 0.19 | 5.14 |
| 125 | 257.28 | 0.29 | 3.49 |
| 126 | 257.9 | 0.61 | 1.63 |
| 127 | 258.09 | 0.2 | 5.09 |
| 128 | 258.36 | 0.26 | 3.8 |
| 129 | 258.56 | 0.2 | 4.89 |
| 130 | 259.06 | 0.5 | 1.99 |
| 131 | 259.3 | 0.24 | 4.17 |
| 132 | 259.51 | 0.21 | 4.81 |
| 133 | 259.71 | 0.2 | 5.07 |
| 134 | 260.19 | 0.48 | 2.07 |
| 135 | 260.31 | 0.12 | 8.67 |
| 136 | 260.38 | 0.07 | 13.76 |
| 137 | 261.04 | 0.66 | 1.51 |

| | | | |
|---|---|---|---|
| 138 | 261.69 | 0.65 | 1.54 |
| 139 | 262.24 | 0.55 | 1.82 |
| 140 | 278.45 | 16.21 | 0.06 |
| 141 | 278.49 | 0.04 | 25.45 |
| 142 | 278.54 | 0.04 | 23.2 |
| 143 | 278.55 | 0.01 | 71.43 |
| 144 | 278.97 | 0.42 | 2.36 |
| 145 | 279.11 | 0.14 | 7.37 |
| 146 | 279.33 | 0.22 | 4.47 |
| 147 | 279.84 | 0.51 | 1.96 |
| 148 | 280.54 | 0.7 | 1.43 |
| 149 | 280.72 | 0.17 | 5.79 |
| 150 | 281.57 | 0.86 | 1.17 |
| 151 | 281.88 | 0.3 | 3.31 |
| 152 | 282.17 | 0.29 | 3.45 |
| 153 | 294.56 | 12.39 | 0.08 |
| 154 | 295.23 | 0.67 | 1.49 |
| 155 | 296.27 | 1.04 | 0.96 |
| 156 | 296.78 | 0.51 | 1.95 |
| 157 | 297.16 | 0.38 | 2.67 |
| 158 | 297.74 | 0.58 | 1.72 |
| 159 | 298.29 | 0.55 | 1.82 |
| 160 | 302.55 | 4.25 | 0.24 |
| 161 | 303.0 | 0.45 | 2.21 |
| 162 | 303.59 | 0.59 | 1.68 |
| 163 | 303.94 | 0.35 | 2.89 |
| 164 | 304.95 | 1.02 | 0.98 |
| 165 | 305.71 | 0.76 | 1.32 |
| 166 | 306.19 | 0.47 | 2.11 |
| 167 | 326.56 | 20.37 | 0.05 |
| 168 | 327.04 | 0.48 | 2.06 |
| 169 | 328.12 | 1.08 | 0.93 |
| 170 | 329.43 | 1.31 | 0.76 |
| 171 | 329.88 | 0.45 | 2.23 |
| 172 | 350.55 | 20.67 | 0.05 |
| 173 | 351.73 | 1.18 | 0.85 |
| 174 | 351.97 | 0.24 | 4.14 |
| 175 | 354.03 | 2.06 | 0.49 |
| 176 | 370.55 | 16.52 | 0.06 |
| 177 | 371.47 | 0.91 | 1.09 |
| 178 | 372.23 | 0.76 | 1.31 |
| 179 | 373.17 | 0.94 | 1.06 |
| 180 | 373.24 | 0.07 | 14.6 |
| 181 | 410.55 | 37.31 | 0.03 |
| 182 | 411.2 | 0.64 | 1.55 |
| 183 | 412.14 | 0.94 | 1.07 |
| 184 | 412.75 | 0.61 | 1.64 |
| 185 | 414.11 | 1.36 | 0.74 |
| 186 | 422.55 | 8.45 | 0.12 |
| 187 | 423.31 | 0.76 | 1.32 |
| 188 | 425.46 | 2.15 | 0.47 |
| 189 | 425.97 | 0.51 | 1.96 |

**Table S8. Spike characteristics for dataset AC detected and sorted using our pipeline.**
This table presents detailed spike information for dataset AC, after sorting with our pipeline. Columns include detected and sorted spike ID, spike timestamp, inter-spike interval (ISI), and instantaneous firing frequency (Hz).

| Spike ID | Spike Timestamp | Inter-spike-interval (ISI) | Instantaneous Frequency (Hz) |
|---|---|---|---|
| 0 | 130.55 | | |
| 1 | 130.62 | 0.08 | 12.94 |
| 2 | 130.76 | 0.14 | 7.31 |
| 3 | 130.82 | 0.06 | 15.87 |
| 4 | 130.88 | 0.06 | 17.3 |
| 5 | 130.94 | 0.05 | 18.76 |
| 6 | 131.01 | 0.07 | 14.08 |
| 7 | 131.06 | 0.05 | 18.32 |
| 8 | 131.11 | 0.05 | 21.14 |
| 9 | 131.16 | 0.05 | 20.45 |
| 10 | 131.21 | 0.06 | 17.99 |
| 11 | 131.27 | 0.06 | 17.27 |
| 12 | 131.33 | 0.06 | 17.73 |
| 13 | 131.37 | 0.05 | 21.6 |
| 14 | 131.42 | 0.05 | 21.69 |
| 15 | 131.48 | 0.06 | 15.72 |
| 16 | 131.55 | 0.06 | 16.16 |
| 17 | 131.61 | 0.06 | 15.67 |
| 18 | 131.69 | 0.08 | 12.85 |
| 19 | 131.74 | 0.05 | 19.88 |
| 20 | 131.81 | 0.08 | 13.14 |
| 21 | 131.89 | 0.08 | 13.3 |
| 22 | 131.96 | 0.07 | 14.31 |
| 23 | 132.05 | 0.09 | 11.04 |
| 24 | 132.14 | 0.09 | 11.59 |
| 25 | 132.24 | 0.1 | 9.55 |
| 26 | 132.32 | 0.08 | 13.28 |
| 27 | 132.45 | 0.13 | 7.68 |
| 28 | 132.54 | 0.1 | 10.29 |
| 29 | 132.62 | 0.08 | 12.84 |
| 30 | 132.7 | 0.08 | 12.17 |
| 31 | 132.87 | 0.16 | 6.06 |
| 32 | 133.0 | 0.13 | 7.43 |
| 33 | 133.1 | 0.1 | 10.16 |
| 34 | 133.21 | 0.11 | 9.03 |
| 35 | 133.46 | 0.24 | 4.08 |
| 36 | 133.73 | 0.28 | 3.63 |
| 37 | 133.97 | 0.24 | 4.12 |
| 38 | 134.21 | 0.24 | 4.16 |
| 39 | 146.56 | 12.34 | 0.08 |
| 40 | 146.86 | 0.3 | 3.31 |
| 41 | 146.95 | 0.09 | 10.88 |
| 42 | 147.07 | 0.12 | 8.65 |
| 43 | 147.31 | 0.24 | 4.11 |
| 44 | 147.53 | 0.22 | 4.57 |
| 45 | 147.71 | 0.19 | 5.4 |
| 46 | 147.95 | 0.23 | 4.26 |
| 47 | 148.14 | 0.19 | 5.25 |
| 48 | 148.26 | 0.12 | 8.18 |
| 49 | 148.65 | 0.39 | 2.56 |
| 50 | 148.95 | 0.3 | 3.38 |
| 51 | 149.14 | 0.19 | 5.15 |
| 52 | 149.32 | 0.18 | 5.62 |
| 53 | 149.55 | 0.23 | 4.36 |
| 54 | 149.84 | 0.3 | 3.39 |
| 55 | 150.21 | 0.37 | 2.72 |

| | | | |
|---|---|---|---|
| 56 | 162.56 | 12.35 | 0.08 |
| 57 | 163.84 | 1.28 | 0.78 |
| 58 | 165.12 | 1.28 | 0.78 |
| 59 | 165.88 | 0.76 | 1.31 |
| 60 | 166.01 | 0.12 | 8.29 |
| 61 | 166.28 | 0.27 | 3.7 |
| 62 | 166.56 | 0.29 | 3.5 |
| 63 | 166.7 | 0.14 | 6.99 |
| 64 | 166.8 | 0.1 | 10.27 |
| 65 | 166.96 | 0.16 | 6.28 |
| 66 | 167.11 | 0.15 | 6.84 |
| 67 | 167.23 | 0.12 | 8.34 |
| 68 | 167.37 | 0.15 | 6.8 |
| 69 | 167.52 | 0.14 | 6.91 |
| 70 | 167.64 | 0.12 | 8.24 |
| 71 | 167.86 | 0.22 | 4.55 |
| 72 | 167.98 | 0.12 | 8.29 |
| 73 | 168.1 | 0.12 | 8.2 |
| 74 | 168.43 | 0.32 | 3.09 |
| 75 | 168.53 | 0.11 | 9.49 |
| 76 | 168.88 | 0.35 | 2.84 |
| 77 | 169.43 | 0.55 | 1.82 |
| 78 | 169.69 | 0.26 | 3.91 |
| 79 | 169.98 | 0.29 | 3.46 |
| 80 | 194.45 | 24.47 | 0.04 |
| 81 | 194.52 | 0.07 | 14.31 |
| 82 | 194.66 | 0.14 | 7.07 |
| 83 | 194.77 | 0.11 | 9.19 |
| 84 | 194.93 | 0.15 | 6.51 |
| 85 | 195.03 | 0.11 | 9.23 |
| 86 | 195.14 | 0.11 | 9.12 |
| 87 | 195.28 | 0.14 | 7.17 |
| 88 | 195.57 | 0.29 | 3.45 |
| 89 | 195.86 | 0.29 | 3.47 |
| 90 | 195.96 | 0.09 | 10.56 |
| 91 | 196.57 | 0.61 | 1.63 |
| 92 | 196.57 | 0.0 | 476.19 |
| 93 | 197.26 | 0.69 | 1.45 |
| 94 | 226.5 | 29.24 | 0.03 |
| 95 | 226.56 | 0.06 | 16.69 |
| 96 | 226.7 | 0.14 | 7.21 |
| 97 | 226.8 | 0.11 | 9.33 |
| 98 | 226.95 | 0.15 | 6.73 |
| 99 | 227.06 | 0.11 | 8.94 |
| 100 | 227.17 | 0.1 | 9.74 |
| 101 | 227.3 | 0.14 | 7.3 |
| 102 | 227.41 | 0.11 | 9.51 |
| 103 | 227.54 | 0.13 | 7.56 |
| 104 | 227.62 | 0.08 | 12.3 |
| 105 | 227.81 | 0.19 | 5.34 |
| 106 | 227.96 | 0.15 | 6.82 |
| 107 | 227.96 | 0.0 | 270.27 |
| 108 | 228.09 | 0.13 | 7.7 |
| 109 | 228.27 | 0.18 | 5.46 |
| 110 | 228.4 | 0.13 | 7.76 |
| 111 | 228.57 | 0.17 | 5.97 |
| 112 | 228.7 | 0.13 | 7.63 |
| 113 | 228.83 | 0.13 | 7.92 |

| | | | |
|---|---|---|---|
| 114 | 229.06 | 0.23 | 4.27 |
| 115 | 254.56 | 25.5 | 0.04 |
| 116 | 254.93 | 0.37 | 2.71 |
| 117 | 255.03 | 0.1 | 9.72 |
| 118 | 255.5 | 0.47 | 2.15 |
| 119 | 256.14 | 0.64 | 1.55 |
| 120 | 256.44 | 0.3 | 3.31 |
| 121 | 257.0 | 0.55 | 1.81 |
| 122 | 257.28 | 0.29 | 3.5 |
| 123 | 257.9 | 0.61 | 1.63 |
| 124 | 258.09 | 0.2 | 5.07 |
| 125 | 258.57 | 0.47 | 2.12 |
| 126 | 259.07 | 0.5 | 2.0 |
| 127 | 259.31 | 0.24 | 4.16 |
| 128 | 259.51 | 0.21 | 4.82 |
| 129 | 259.71 | 0.2 | 5.07 |
| 130 | 260.19 | 0.48 | 2.07 |
| 131 | 261.04 | 0.85 | 1.18 |
| 132 | 261.69 | 0.65 | 1.54 |
| 133 | 262.24 | 0.55 | 1.82 |
| 134 | 278.5 | 16.26 | 0.06 |
| 135 | 279.11 | 0.62 | 1.62 |
| 136 | 279.34 | 0.22 | 4.48 |
| 137 | 279.85 | 0.51 | 1.96 |
| 138 | 280.55 | 0.7 | 1.43 |
| 139 | 280.72 | 0.17 | 5.81 |
| 140 | 281.58 | 0.85 | 1.17 |
| 141 | 281.88 | 0.3 | 3.3 |
| 142 | 282.17 | 0.29 | 3.45 |
| 143 | 294.56 | 12.4 | 0.08 |
| 144 | 295.23 | 0.67 | 1.49 |
| 145 | 296.27 | 1.04 | 0.96 |
| 146 | 296.79 | 0.51 | 1.95 |
| 147 | 297.16 | 0.37 | 2.67 |
| 148 | 297.74 | 0.58 | 1.72 |
| 149 | 298.29 | 0.55 | 1.82 |
| 150 | 302.55 | 4.26 | 0.23 |
| 151 | 303.0 | 0.45 | 2.21 |
| 152 | 303.6 | 0.59 | 1.68 |
| 153 | 303.94 | 0.34 | 2.9 |
| 154 | 304.96 | 1.02 | 0.98 |
| 155 | 305.72 | 0.76 | 1.32 |
| 156 | 306.19 | 0.47 | 2.12 |
| 157 | 326.56 | 20.38 | 0.05 |
| 158 | 327.05 | 0.48 | 2.07 |
| 159 | 328.13 | 1.08 | 0.93 |
| 160 | 329.44 | 1.31 | 0.76 |
| 161 | 329.88 | 0.45 | 2.23 |
| 162 | 351.74 | 21.85 | 0.05 |
| 163 | 351.98 | 0.24 | 4.17 |
| 164 | 354.03 | 2.06 | 0.49 |
| 165 | 370.56 | 16.53 | 0.06 |
| 166 | 371.47 | 0.91 | 1.1 |
| 167 | 372.23 | 0.76 | 1.31 |
| 168 | 373.24 | 1.01 | 0.99 |
| 169 | 410.56 | 37.32 | 0.03 |
| 170 | 412.14 | 1.58 | 0.63 |
| 171 | 412.75 | 0.61 | 1.64 |

| | | | |
|---|---|---|---|
| 172 | 414.11 | 1.36 | 0.74 |
| 173 | 422.56 | 8.45 | 0.12 |
| 174 | 423.31 | 0.75 | 1.33 |
| 175 | 425.46 | 2.15 | 0.47 |
| 176 | 425.97 | 0.51 | 1.96 |
